# Live-cell imaging and CLEM reveal the existence of ACTN4-dependent ruffle-edge lamellipodia acting as a novel mode of cell migration

**DOI:** 10.1101/2024.05.20.594911

**Authors:** Haruka Morishita, Katsuhisa Kawai, Youhei Egami, Kazufumi Honda, Nobukazu Araki

## Abstract

α-Actinin-4 (ACTN4) expression levels are correlated with the invasive and metastatic potential of cancer cells; however, the underlying mechanism remains unclear. Here, we identified ACTN4-localized ruffle-edge lamellipodia using live-cell imaging and correlative light and electron microscopy (CLEM). BSC-1 cells expressing EGFP-ACTN4 showed that ACTN4 was most abundant in the leading edges of lamellipodia, although it was also present in stress fibers and focal adhesions. ACTN4 localization in lamellipodia was markedly diminished by phosphoinositide 3-kinase inhibition, whereas its localization in stress fibers and focal adhesions remained. Furthermore, overexpression of ACTN4, but not ACTN1, promoted lamellipodial formation. Live-cell analysis demonstrated that ACTN4-enriched lamellipodia are highly dynamic and associated with cell migration. CLEM revealed that ACTN4-enriched lamellipodia exhibit a characteristic morphology of multilayered ruffle-edges that differs from canonical flat lamellipodia. Similar ruffle-edge lamellipodia were observed in A549 and MBA-MD-231 invasive cancer cells. ACTN4 knockdown suppressed the formation of ruffle-edge lamellipodia and cell migration during wound healing in A549 monolayer cultures. Additionally, membrane-type 1 matrix metalloproteinase was observed in the membrane ruffles, suggesting that ruffle-edge lamellipodia have the ability to degrade the extracellular matrix and may contribute to active cell migration/invasion in certain cancer cell types.

## Introduction

Cell migration is essential for many physiological and pathological processes, such as embryonic development, wound healing, inflammation, and cancer cell invasion. Although diverse modes of cell migration, including membrane blebbing and cytoplasmic streaming, are used for various purposes by different cell types [1], the migration mechanism through lamellipodial protrusions appears to be the primary one in many cases. In general, lamellipodia are defined as thin, sheet-like cell protrusions that constitute the actin cytoskeleton-based motile apparatus, which is indispensable for promoting the movement of migrating cells [2]. Cell migration is established by progressive extension of lamellipodia using basal cell adhesion as a scaffold and retraction of the posterior tail of the cell [3,4]. The extension of lamellipodia is initiated by the formation of a network of branched actin filaments (F-actin) through the Arp2/3 complex, which is activated by nucleation promoting factors such as the WAVE regulatory complex (WRC)[5–8]. The Rho-family GTPase Rac1 is a major switch molecule that regulates lamellipodial formation [9]. GTP-bound Rac1, which is targeted to the plasma membrane, activates the WRC-Arp2/3-F-actin nucleation axis, thereby facilitating membrane protrusion [10,11]. Dynamic changes in the actin cytoskeleton architecture inside the formed lamellipodia are driven by a variety of actin-binding proteins, including lamellipodin [12], cortactin [13], cofilin [14], α-actinin, and myosins [15]. These proteins are known to be regulated by membrane phospholipids such as phosphatidylinositol 4,5-bisphosphate (PI(4,5)P_2_) and phosphatidylinositol 3,4,5-triphosphate (PI(3,4,5)P_3_) [16]. PI(3,4,5)P_3_ is produced from PI(4,5)P_2_ by class I phosphoinositide 3-kinase (PI3K) and is concentrated in projecting lamellipodia, as well as GTP-bound Rac1 [10,17–19].

When lamellipodia detach from the substratum and roll up, their structure and motion are termed membrane ruffles and ruffling, respectively. Membrane ruffles arising from the cell periphery move centripetally to the dorsal cell surface. Although the term ‘membrane ruffles’ is commonly used for various structures, including peripheral ruffles [18,20], circular ruffles (macropinocytic cups) [21,22], and dorsal circular ruffles [23], the morphology and functions of membrane ruffles may vary considerably depending on the cell type and specific circumstances. α-Actinin (ACTN) is an actin-binding protein, and its two isoforms, ACTN1 and ACTN4, exist in non-muscle cells. ACTN1 primarily localizes to focal adhesion plaques and anchors F-actin stress fibers to the plasma membrane with proteins, including talin, vinculin, and integrin [24,25]. In 1998, ACTN4 was cloned as a new isoform associated with cancer cell invasion [26]. Although ACTN1 and ACTN4 are highly homologous in their nucleotide and amino acid sequences, ACTN4 is presumed to be preferentially involved in more dynamic cellular structures, including lamellipodia and membrane ruffles [22,26– 28]. In recent years, *in vitro* and *in vivo* studies have demonstrated a correlation between increased ACTN4 expression and several advanced cancers including oral, breast, colorectal, and pancreatic cancers [29– 31]. Several lines of evidence have implicated ACTN4 in the invasion and metastasis of cancer cells [30,32– 34]. Evidence now describes ACTN4 as a useful biomarker indicative of cancer progression and prognosis, and a potential therapeutic target in patients with various cancers [29,32]. However, the precise role of ACTN4 in lamellipodial formation associated with cell migration remains to be elucidated.

Recently, we found highly motile lamellipodia with DENND1B-positive structures at the bottom, which were most conspicuously observed in BSC-1 cells [35]. In exploring key molecules characterizing the lamellipodia, we found that ACTN4 is abundantly observed in the lamellipodia. In this study, live-cell imaging and correlative light-electron microscopy (CLEM) demonstrate that ACTN4-enriched lamellipodia exhibit special morphological and functional phenotypes of cell migration and invasion. We further show that the formation of a special form of lamellipodia is highly dependent on PI3K activity. Importantly, ACTN4 knockdown inhibits lamellipodia formation and cell migration. Since the leading edge of lamellipodia shows extracellular matrix degradation activity, the formation of ACTN4-enriched lamellipodia may act as a novel mode of cell migration and invasion. Our findings provide new insights into why ACTN4 is associated with enhanced cell migration and may be linked to the progression of certain types of cancer.

## Materials and Methods

### Reagents and Plasmids

Full-length cDNA encoding human ACTN4 was amplified by PCR. The fragment was cloned into the pEGFP vector. pmCherry-Lifeact was constructed by cloning the synthesized cDNA fragment of Lifeact into pmCherry vector. pEGFP-ACTN1 (Addgene #11908) was obtained from Dr. Carol Otey (UNC-Chapel Hill) and pTriEx/mCherry-PA-Rac1 (Addgene #22024) was obtained from Dr. Klaus Hahn (UNC-Chapel Hill). pmCitrine-Rac1, pmCitrine-Rac1Q61L, pmCitrine-Rac1T17N, pCFP-membrane, pmCitrine-Akt-PH, and pYFP-Btk-PH domains were gifts from Dr. Joel A. Swanson (University of Michigan). The pmCherry-Akt-PH domain and pmCherry-membrane were generated by replacing mCitrine or CFP cDNA with mCherry cDNA. Other reagents were purchased from FUJIFILM Chemicals (Osaka, Japan) or Nacalai Tesque (Kyoto, Japan), unless otherwise indicated.

### Cell culture, and transfection

BSC-1 cells, epithelial cells of African green monkey kidney origin, were obtained from the JCRB Cell Bank (Osaka, Japan). HeLa, MDCK-II, B16, A549, A431, and RAW264 cells were obtained from the Riken Cell Bank (Tsukuba, Japan). MDA-MB-231 cells were obtained from the ATCC (Manassas, VA). The cells were cultured in Dulbecco’s modified Eagle’s medium (DMEM) supplemented with 10% heat-inactivated fetal bovine serum (FBS) and antibiotics (penicillin and streptomycin) at 37°C in a humidified atmosphere containing 5% CO_2_. Cells were transfected with plasmids using the Neon Transfection System (Life Technologies, Carlsbad, CA), according to the manufacturer’s protocols. The cells were seeded onto 13-or 25-mm circular coverslips in culture dishes containing the culture media. The cells were used for experiments 12–24 h after transfection.

### ACTN4 siRNA knockdown and wound healing assay

ACTN4 knockdown experiments were performed using Stealth Select RNAi (Thermo Fisher Scientific). A549 cells were transfected with small interfering RNA (siRNA) targeting human ACTN4 (#1 HSS100124, #2 HSS100125, #3 HSS188682), human ACTN1 (HSS100130), or negative control siRNA (Med GC Duplex#3) using Lipofectamine RNAiMAX (Thermo Fisher Scientific) according to the manufacturer’s instructions. The cells were subjected to experiments 48–72 h after transfection. The knockdown effects of ACTN1 and ACTN4 siRNAs were confirmed by immunoblotting and immunofluorescence assays. We quantified the knockdown efficiency by measuring the fluorescence intensity of ACTN4 immunofluorescence staining in A549 cells using the MetaMorph imaging system. Each group of cells was immunostained and imaged under identical conditions. The fluorescence intensity relative to that of the negative control siRNA-treated cells was calculated after background subtraction.

To observe migrating cells, a confluent monolayer of A459 cells on coverslips was scratched with a 200 µL pipette tip. After 5–24 h of incubation in the presence of 50 nM phorbol 12-myristate 13-acetate (PMA), the cells that migrated from the edge of the artificial wound were observed. A wound healing assay was performed to determine the cell migration capacity by measuring the changes in wound width by subtracting the width of the wounds at 5 h from that at 0 h after scraching.

### Immunoblotting

Total protein lysates were prepared with a lysis buffer consisting of 50 mM Tris-HCl (pH 7.5), 150 mM NaCl, 1% Triton X-100, 0.5% sodium deoxycholate, 0.1% sodium dodecyl sulfate (SDS), 0.1% CHAPS, and protease inhibitor cocktail (Nacalai Tesque, Kyoto, Japan). Equal amounts of protein were denatured and reduced with sample buffer containing 1% SDS and 2.5% 2-mercaptoethanol. The samples were subjected to 10% SDS-PAGE and transferred to polyvinylidene difluoride membranes (Millipore, Bedford, MA, USA). The membranes were immunoreacted with anti-ACTN4 mouse monoclonal IgG (Abnova, clone 13G9, dilution 1:10000), or anti-ACTN1 rabbit polyclonal IgG (Abcepta, clone RB21901, dilution 1:1000). Anti-GAPDH mouse monoclonal antibody (Ambion, clone 6C5, dilution 1:10000) was used to verify equal amounts of total protein loading. After washing, the membranes were incubated with HRP-conjugated anti-rabbit IgG or anti-mouse IgG secondary antibodies and developed using the ECL Prime Western Blotting Detection System (Cytiva, Tokyo, Japan).

### Live-cell imaging

For live-cell imaging, cells were cultured on 25-mm circular coverslips. Before imaging, the culture medium was replaced with Ringer’s buffer (RB) consisting of 155 mM NaCl, 5 mM KCl, 1 mM MgCl_2_, 2 mM Na_2_HPO_4_, 10 mM glucose, 10 mM HEPES and 0.5 mg/ml bovine serum albumin (BSA) at pH 7.2. The 25-mm circular coverslip was assembled in an RB-filled chamber (Attofluor, #A7816, Molecular Probes) and placed on a 37°C thermo-controlled stage in Leica DMI6000B inverted microscope (Leica Microsystems, Wetzlar, Germany). Live-cell images were acquired with an oil-immersion 100× objective lens (HC PL Fluotar PH3, NA=1.32 or HCX PL Apo, NA=1.4) using an Orca Flash 4.2 sCMOS camera (Hamamatsu Photonics, Shizuoka, Japan) under the control of the MetaMorph Imaging software (Molecular Devices). Kymographs were generated using MetaMorph software by taking 1-pixel wide rectangular regions in the direction of lamellipodial extension.

### Optogenetics of photoactivatable Rac1 under the fluorescence microscope

Cells transfected with pTriEx/mCherry-PA-Rac1Q61L and pEGFP-ACTN4 were cultured on 25 mm circular coverslips for 16-24 h. The coverslips were subjected to live-cell imaging as described above. Optogenetic control of photoactivatable (PA)-Rac1 under the Leica DMI6000B microscope was achieved using a journal (macro-program) of MetaMorph software, which automates the acquisition of time-lapse images and photoactivation during the image acquisition interval [36]. Because the light oxygen voltage 2 (LOV2) domain of PA-Rac1 is a photosensor for blue light (360–500 nm), we activated PA-Rac1 by illuminating the cells through a cyan fluorescent protein (CFP) excitation filter (Chroma ET436/24 nm). When illuminated with blue light, the LOV2 domain undergoes a conformational change that enables downstream effector binding to Rac1, indicating that the cells exhibit a constitutively active Rac1 phenotype [37,38].

### Immunofluorescence microscopy

BSC-1 cells on coverslips were fixed with 4% paraformaldehyde (PFA) in 0.1M phosphate buffer (pH 7.4) containing 6% sucrose for 30 min at room temperature. After rinsing with phosphate-buffered saline (PBS), the cells were permeabilized with 0.25% Triton X-100/PBS and blocked with 1% BSA/0.5% gelatin/PBS for 10 min each. The cells were then incubated with rabbit polyclonal anti-ACTN4 antibody (dilution 1:100)[26] or mouse monoclonal anti-ACTN4 antibody (Abnova, Clone 13G9, dilution 1:300) for 60 min, followed by incubation with a secondary antibody conjugated with Alexa Fluor 488 or 594 (Molecular Probes, dilution 1:1000). An anti-vinculin mouse monoclonal antibody (V9264; Sigma-Aldrich, dilution 1:200) was used as a focal adhesion marker. Membrane-type 1 matrix metalloproteinase (MT1-MMP; also known as MMP14) was detected in A549 cells fixed with 2% paraformaldehyde for 15 min using anti-MT1-MMP mouse IgM monoclonal antibody (AM1832a, Abcepta, dilution 1:100)), followed by Alexa Fluor 568-conjugated anti-mouse IgM (Molecular Probes). F-actin was stained with Alexa Fluor 350-or 568-conjugated phalloidin (Molecular Probes) or Acti-stain 670 phalloidin (Cytoskeleton Inc.). Nuclear staining was performed using 4’,6-diamidino-2-phenylindole (DAPI) solution (1 μg/ml in PBS).

The specimens were mounted onto glass slides and examined under a confocal microscope (LSM710, Carl Zeiss, Oberkochen, Germany) or an epifluorescence microscope (Leica DMI6000B, Germany). Some specimens were observed by a confocal-based super-resolution microscope (Nikon AX with NSPARC detector, Japan) using a plan apo 60× NA=1.42 objective lens.

### Classification and quantitation of cells based on ACTNs localization patterns

BSC-1 cells immunostained for ACTN4 and expressing EGFP-ACTN4 or ACTN1 were classified into three types according to ACTNs localization patterns, as shown in Fig. 1 and Fig. 2. Lamellipodia localization type (L-type) cells were defined as cells in which ACTNs intensively localize in lamellipodial leading edges but not in stress fibers or focal adhedions. By phase-contrast microscopy, the leading edge of lamellipodia in L-type cells was characterized as phase-dark. Cells in which ACTNs predominantly localize in stress fibers and focal adhesions were defined as stress fiber/focal adhesion localization type (SF-type). SF-type cells do not have lamellipodia. Cells in which ACTNs moderately localize in both lamellipodia and stress fibers/focal adhesions were defined as intermediate type (I-type). Lamellipodia in I-type cells do not have phase-dark structures in their leading edges. The proportion of the three cell types in each BSC-1 cell group was quantified by counting ≥50 cells on each coverslip under a microscope. The data are presented as the mean ± standard error of the mean (SEM) of n≥6 coverslips from three independent experiments.

**Figure 1.**
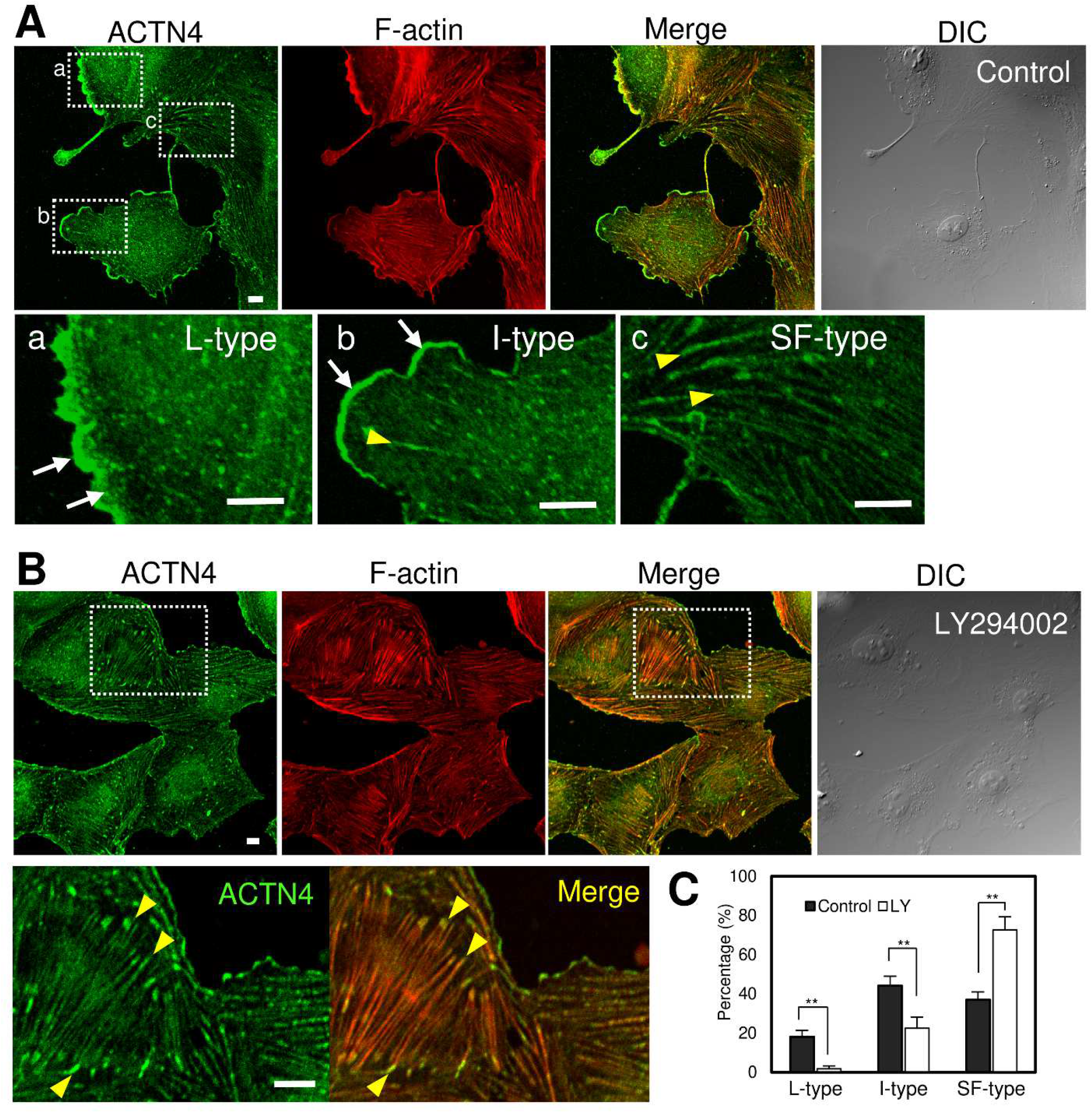
Confocal microscopy showing endogenous ACTN4 localization and F-actin distribution in control BSC-1 cells (A) and LY294002-treated cells (B). Cells were fixed and immunostained with an anti-ACTN4 antibody, followed by an Alexa Fluor 488-labeled secondary antibody. F-actin was detected using Alexa Fluor 568-phalloidin. (A) In control cells, ACTN4 was localized to lamellipodia (arrows), actin stress fibers, and focal adhesions (arrowheads). The three boxed areas (a, b, and c in the ACTN4 image) are magnified in the lower panel. (B) In cells pretreated with 25 μM LY294002 for 20 min, ACTN4-enriched lamellipodia were scarcely observed, although ACTN4 localization in stress fibers and focal adhesions was evident. Magnified images of the boxed areas are shown below. DIC, differential interference contrast images. Scale bars, 10 µm. (C) The percentage of cells showing three ACTN4 localization types: lamellipodia localization (L)-type, intermediate (I)-type, and stress fiber/focal adhesion localization (SF)-type, was calculated by counting >60 cells on each coverslip for the control (gray bars) and 25 μM LY294002 (LY)-treated cells (open bars). Each bar represents the mean ± SEM of n=6 coverslips from three independent experiments. \*\**P*<0.01.

**Figure 2.**
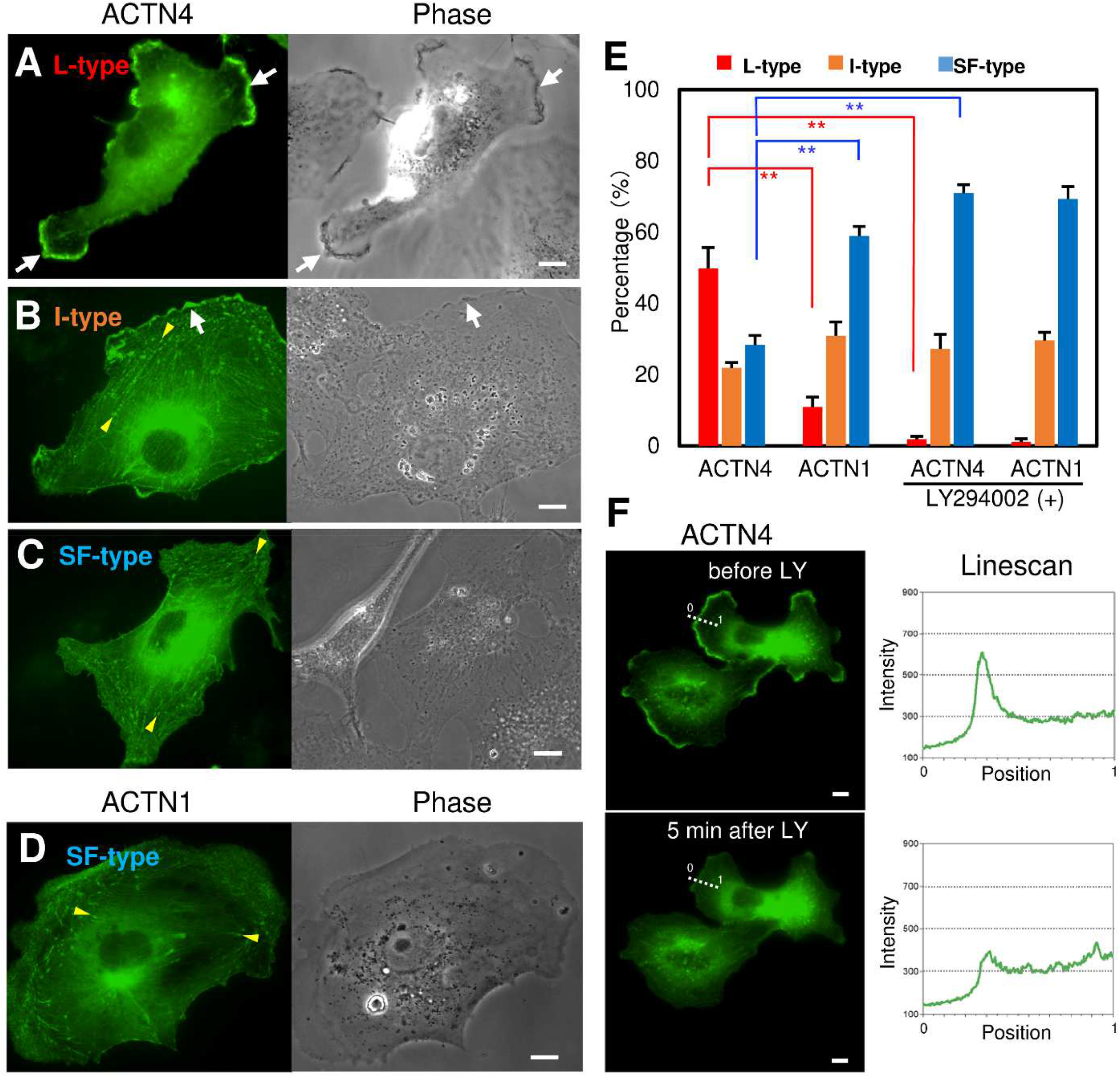
Overexpression of ACTN4, but not ACTN1, enhances ACTN-enriched lamellipodial formation in a PI3K-dependent manner. Based on the ACTN localization pattern, BSC-1 cells transfected with pEGFP-ACTN4 were classified into three types: L-type (A), I-type (B), and SF-type (C). Among EGFP-ACTN1 overexpressing cells, SF-type cells were the most abundant (D). Arrows indicate lamellipodia. Arrowheads point focal adhesions.(E) Percentages of cells showing three ACTN localization types, L-type (red bars), I-type (orange), and SF-type (blue), in ACTN4-or ACTN1-overexpressing cells before and after treatment with 25 μM LY294002 (LY). \*\**P*<0.01, n=6. (F) EGFP-ACTN4 in the lamellipodium was markedly diminished after the addition of LY294002. Linescan analysis (right panel) along the dotted line in the left images shows a decrease in the fluorescence intensity of EGFP-ACTN4 in lamellipodia after the addition of LY294002. The corresponding video is available in the supplementary materials (Video 1). All scale bars, 10 µm.

Cells in contact with the surrounding cells all around and without free cell edges were excluded from counting because they could not extend lamellipodia. In addition, cells showing abnormal EGFP-fusion protein aggregation, probably owing to exstream overexpression of the EGFP-fusion proteins, were excluded.

### Correlative light-electron microscopy (CLEM)

Cells transfected with pEGFP-ACTN4 were cultured on grided coverslips in 35 mm dishes and fixed with 2% glutaraldehyde in 0.1M phosphate buffer (pH 7.4) for 30 min. Coverslips in a PBS-filled dish were observed under a Zeiss LSM710 confocal laser microscope using a 40x Plan Apochromat water immersion objective lens (N.A. 1.0). EGFP-ACTN4 fluorescence and DIC images were obtained using ZEN software. The coverslips were then post-fixed with 1% OsO_4_ and processed for scanning electron microscopy as previously described [39]. Critical point-dried specimens were coated with osmium using an osmium plasma coater (OPC40, Nippon Laser & Electronics, Japan) and observed with a field-emission scanning electron microscope (EM) (Hitachi S900, Japan) at 6 kV. The same cell was photographed through the confocal laser microscope was searched and photographed using scanning EM. Scanning EM and confocal fluorescence images were overlaid using Adobe Photoshop. Similarly, cells immunostained with anti-ACTN4 were processed to scanning EM after fluorescence microscopy for subjecting to CLEM.

### Fluorescein gelatin degradation assay

Sterilized coverslips were coated with 0.1 mg/ml poly-L-lysine for 20 min, washed with PBS, and fixed with 0.5% glutaraldehyde for 10 min. After three washes with PBS, the coverslips were inverted on a 40-μl drop of 0.2% fluorescein (FITC)-labeled gelatin (Thermo Fisher Scientific) in 2% sucrose in PBS and incubated for 10 min. After washing with PBS, free aldehyde groups were quenched with 1 mg/ml sodium borohydride in PBS for 5 min, washed three times in PBS, and incubated with 1 ml of complete medium for 30 min. Cells were plated on fluorescent gelatin-coated coverslips, incubated for 30-60 min, and then fixed with 4% paraformaldehyde for 30 min. The samples were processed for staining with anti-ACTN4 monoclonal antibody and ActiStain 670 phalloidin. The coverslips were imaged using a Nikon AX confocal microscope.

### Statistical analysis

For quantitative analysis, data were obtained from at least three independent measurements and expressed as the mean ± standard error of the mean (SEM). Statistical significance was calculated using a two-tailed Student’s *t*-test or one-way analysis of variance (ANOVA), followed by Tukey’s test. Differences were considered statistically significant at *P* <0.05.

## Results

### Localization of endogenous ACTN4 and its PI3K dependency in BSC-1 cells

First, we examined the localization of endogenous ACTN4 in control BSC-1 cells by anti-ACTN4 immunofluorescence and phalloidin staining for F-actin. Confocal microscopy revealed that ACTN4 is abundantly localized in F-actin-based structures, including lamellipodia, actin stress fibers, and focal adhesions (Fig. 1A). However, the localization pattern of ACTN4 varies considerably among cells. Typically, BSC-1 cells with ACTN4-enriched lamellipodia had fewer stress fibers. In contrast, cells without lamellipodia showed ACTN4-positive stress fibers and focal adhesions. Based on the ACTN4 localization pattern, we classified the cells into three types, examples of which are shown in Fig.1A: (i) lamellipodia type (L-type), in which ACTN4 predominantly localizes in lamellipodia (Fig.1A, a); (ii) intermediate type (I-type), in which ACTN4 moderately localizes in both lamellipodia and stress fibers/focal adhesions (Fig. 1A, b); and (iii) stress fiber type (SF-type), in which ACTN4 predominantly localizes in stress fibers/focal adhesions (Fig. 1A, c). We then calculated the percentage of each type in BSC-1 cells by counting the number of each type among ≥50 cells/coverslip (total 6 coverslips from three independent experiments) under fluorescence microscopy of anti-ACTN4 immunoreaction and phalloidin staining. Among control BSC-1 cells, L-type cells accounted for 18.1 ± 3.2% (n=6), I-type for 44.2 ± 4.8% (n=6), and SF-type for 37.0 ± 4.0% (n=6) (Fig. 1C, grey bars).

Phosphoinositides, such as PI(4,5)P_2_ and PI(3,4,5)P_3_, are known to be crucial regulators of actin cytoskeleton organization and membrane trafficking [16,40,41]. Fraley et al. (2003) reported that PI(3,4,5)P_3_ binds to the N-terminal calponin homology domain of actinin and downregulates its actin-bundling ability [42]. Therefore, we examined the effect of LY294002, a PI3K inhibitor that inhibits the production of PI(3,4,5)P_3_ from PI(4,5)P_2_, on the localization of ACTN4 in BSC-1 cells. ACTN4-enriched lamellipodia were scarcely observed in LY294002-treated cells. The localization of ACTN4 in stress fibers and focal adhesions was not significantly affected by treatment with the PI3K inhibitor LY294002 (Fig. 1B). Counting the number of cells with ACTN4-enriched lamellipodia revealed that LY294002 treatment greatly reduced the proportion of L-type cells instead of increasing the number of SF-type cells (Fig. 1C, open bars).

### Overexpression of ACTN4, but not ACTN1, increases lamellipodia formation

Next, we examined the localization of exogenously overexpressed ACTN4 or ACTN1 in BSC-1 cells by transfecting pEGFP-ACTN4 or pEGFP-ACTN1 together with pmCherry-Lifeact. Although both EGFP-ACTN1 and EGFP-ACTN4 were observed in lamellipodia, stress fibers, and focal adhesions, they showed distinct distribution quantities among these actin-based structures (Fig.2A-D, Supplementary Fig. S1, S2A). In EGFP-ACTN4-overexpressing cells, ACTN4 was highly abundant in lamellipodia (Fig.2A, Supplementary Fig. S1). In contrast, ACTN1 was enriched in stress fibers and focal adhesions at both ends of stress fibers in EGFP-ACTN1-overexpressing cells. ACTN1-enriched lamellipodia were observed less frequently (Fig.2D, Supplementary Fig. S2A). To examine the impact of ACTNs overexpression, we calculated the percentage of each type of cell overexpressing EGFP-ACTN4 or EGFP-ACTN1 by counting the number of each type of cell among ≥ 50 cells/coverslip (total 6 coverslips from three independent experiments) under phase-contrast and fluorescence microscopy of mCherry-Lifeact and EGFP-ACTNs. Among ACTN4-overexpressing cells, L-type cells accounted for ∼50%, I-type for ∼20%, and SF-type for ∼30%. In contrast, among ACTN1-overexpressing cells, L-type accounted for ∼10%, I-type for ∼30%, and SF-type for ∼60% (Fig.2E). Thus, the proportion of L-type cells among ACTN4-overexpressing cells was significantly higher than that among ACTN1-overexpressing cells (*P*<0.01, n=6).

Compared with control BSC-1 cells, of which ∼18% cells exhibited L-type (Fig.1E), overexpression of ACTN4 increased the proportion of L-type cells more than three-fold; in contrast, ACTN1 overexpression decreased L-type cells to ∼10%, and instead increased SF-type cells to ∼60% (Fig. 2E). These results suggest that ACTN4 overexpression may facilitate ACTN4-enriched lamellipodia formation, whereas ACTN1 overexpression may have a higher propensity to form stress fibers and focal adhesions in BSC-1 cells.

### PI3K activity is indispensable for ACTN4 localization to the leading edge of ACTN4-enriched lamellipodia, but not to stress fibers or focal adhesions

Live-cell imaging and high-resolution confocal microscopy of BSC-1 cells expressing EGFP-ACTN4 showed that EGFP-ACTN4 in the lamellipodial leading edge was markedly diminished after the addition of the PI3K inhibitor, whereas EGFP-ACTN4 in stress fibers and focal adhesions appeared to have remained (Fig.2F, Supplementary Fig. S3, Video 1). Quantitation of ACTN4 localization types revealed that the percentage of the L-type was drastically reduced to only 2% by PI3K inhibition, and instead the SF-type increased to ∼70% (Fig.2E). These findings suggest that PI3K activity is indispensable for ACTN4 localization to the leading edge of ACTN4-enriched lamellipodia, whereas that to stress fibers and focal adhesions may not require PI3K activity.

As EGFP-ACTN1 was predominantly observed in stress fibers and focal adhesions in control BSC-1 cells, changes in EGFP-ACTN1 localization after PI3K inhibition were not as evident as in the case of ACTN4 by confocal microscopy (Supplementary Fig. S2). However, the quantitation of ACTN1-overexpressing cells showed that the percentage of L-type cells significantly decreased after PI3K inhibition (Fig. 2E). Upon PI3K inhibition, ACTN1-and ACTN4-overexpressing cells showed similar ACTNs localization patterns, in which only a small percentage of cells exhibiting L-type.

Because PI(3,4,5)P_3_, a major product of class I PI3K, is known to bind to the calponin homology domain 2 (CH2) of ACTN molecules [43,44], it is possible that increased levels of PI(3,4,5)P_3_ in the plasma membrane recruit ACTN4 to the leading edge of lamellipodia. Therefore, we visualized PI(3,4,5)P_3_/PI(3,4)P_2_ by co-expressing mCherry-Akt pleckstrin homology domain (Akt-PH) and EGFP-ACTN4 in live BSC-1 cells. As expected, mCherry-Akt-PH signal levels were particularly high in the region of extending lamellipodia enriched with EGFP-ACTN4 (Fig. 3A). However, mCherry-Akt-PH signals were scarcely observed in stress fibers or focal adhesions. When LY294002 was applied to the cells, mCherry-Akt-PH signals in ACTN4-enriched lamellipodia immediately disappeared.

**Figure 3.**
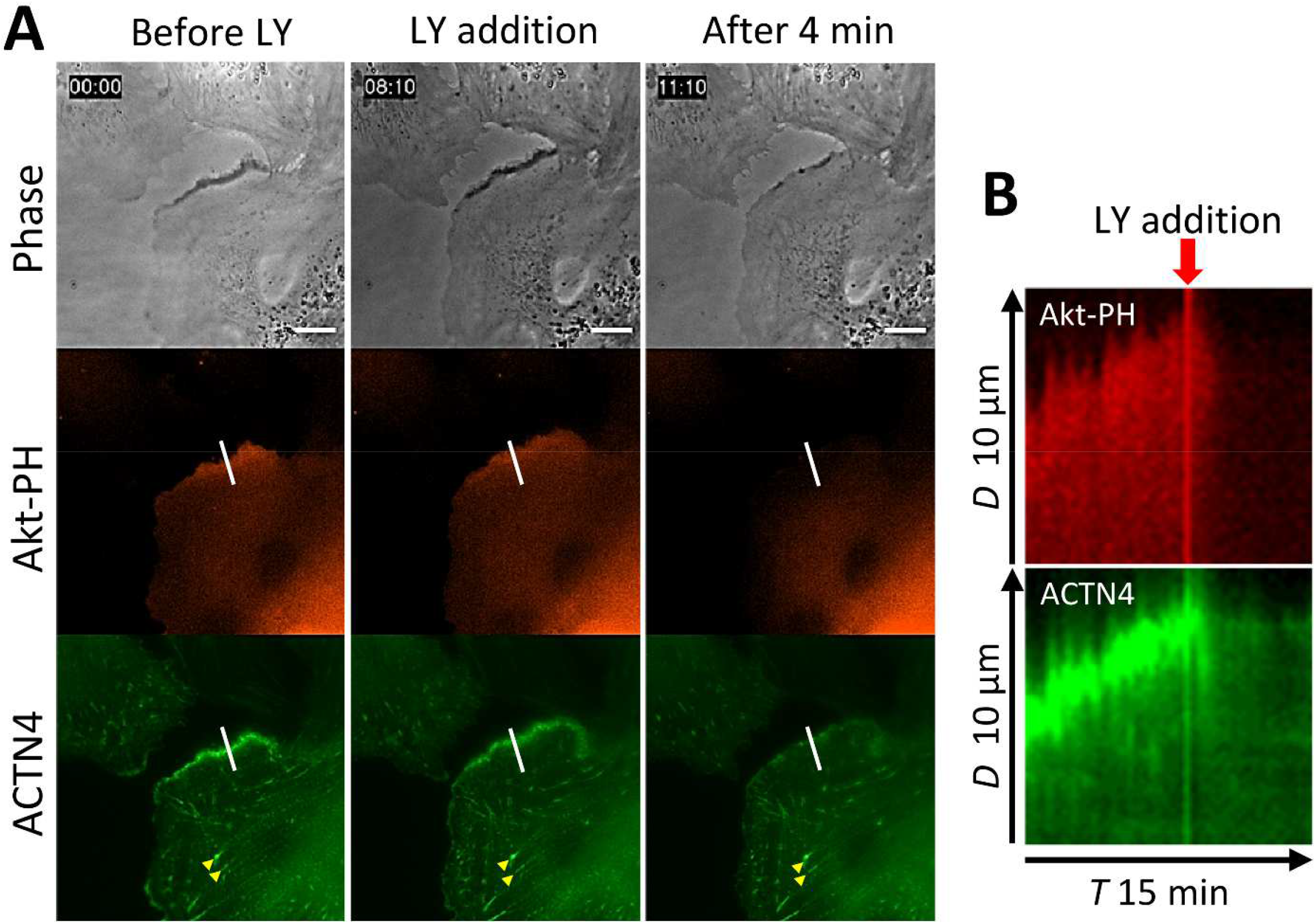
EGFP-ACTN4 was diminished almost simultaneously with mCherry-Akt-PH domain (Akt-PH) monitoring PI(3,4,5)P_3_ and PI(3,4)P_2_ by the addition of LY294002. (A) Time-lapse images of live-BSC-1 cells co-expressing EGFP-ACTN4 and mCherry-Akt-PH. Eight minutes after the start of the time-lapse imaging, 25 μM LY294002 (LY) was added to the cells. Arrowheads indicate that ACTN4 localizes in focal adhesions and stress fibers even after LY addition. The times elapsed from the first frame acquisition are shown in the upper-left corner. (B) Kymographs from the time series along the white line in Akt-PH and ACTN4 images of A. The constructed kymographs depict the trajectory of lamellipodial movement and fluorescent signals. Rows of pixels at the line are laid side-by-side in a montage to create a time (*T*) vs. distance (*D*) plot. Both fluorescent signals of Akt-PH and ACTN4 diminished almost simultaneously within a few minutes of LY addition. The arrow indicates the time point of LY addition. Scale bar, 10 µm.

Almost simultaneously, EGFP-ACTN4 greatly diminished from the leading edge of lamellipodia (Fig. 3A and B), although EGFP-ACTN4 in the stress fibers and focal adhesions remained (arrowheads in Fig. 3A, Supplementary Fig. S3). A decrease in PI(3,4,5)P_3_/PI(3,4)P_2_ concentrations in the membrane of lamellipodia was verified by ratio imaging of mCitrine-Akt-PH/mCherry-membrane (Supplementary Fig. S4A). Using the YFP-Btk-PH probe, which binds only to PI(3,4,5)P_3_, we obtained results similar to those obtained with the Akt-PH probe (Supplementary Fig. S4B, Video 2). These observations support the hypothesis that PI(3,4,5)P_3_ is an important factor in ACTN4 localization to membranes forming ACTN4-enriched lamellipodia.

### ACTN4-enriched lamellipodia are highly dynamic and invasive structures

To characterize the dynamics of ACTN4-enriched lamellipodia, time-lapse movies of live BSC-1 cells expressing EGFP-ACTN4 were created. A movie of EGFP-ACTN4-expressing BSC-1 cells showed that extending lamellipodia were rich in ACTN4, whereas retracting or static lamellipodia-like protrusions were ACTN4-modest (Fig. 4A, Video 3). The moving speed of the ACTN4-enriched lamellipodia was much faster than that of ACTN4-modest lamellipodia. Therefore, the dynamics of lamellipodia were assessed by kymograph analysis using MetaMorph software (Fig. 4B and C). ACTN4-enriched lamellipodia always extended forward at an average speed of 0.37 ± 0.15 µm/min (n=15 lamellipodia collected from 3 independent experiments), whereas ACTN4-modest flat lamellipodia moved with repeated extension and retraction. Because the leading edge of flat lamellipodia fluctuated with almost equal amplitude of extension and retraction, only a small net advance (0.07 ± 0.04 µm/min, n=15, collected from 3 independent experiments) was observed (Fig. 4C).

**Figure 4.**
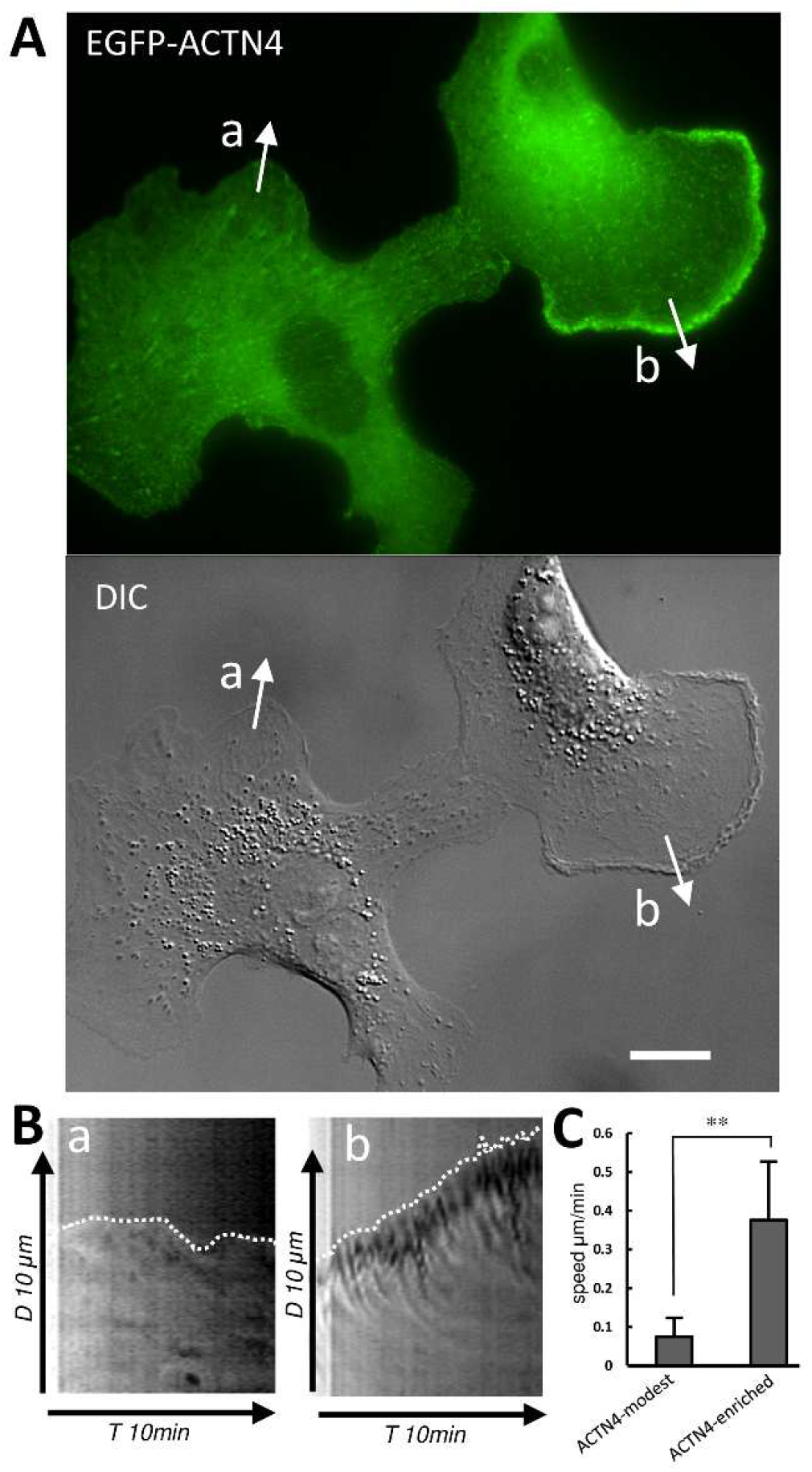
Comparison of the dynamics of ACTN4-enriched and ACTN4-modest lamellipodia in BSC-1 cells. The left cell shows ACTN4-modest lamellipodia (a). The right cell has an ACTN4-enriched lamellipodium, in which the leading edge shows a wavy appearance (b). Scale bar: 10 µm. The corresponding video is available in the supplementary materials (Video 3). **(B)** Kymograph analysis of ACTN4-modest (arrow, a) and ACTN4-enriched lamellipodia (arrow, b). ACTN4-enriched lamellipodia always extended forward, whereas ACTN4-modest flat lamellipodia moved with repeated extension and retraction. (**C**) Quantitation of moving spped of lamellipodial edges based on the kymograph analysis. Comparing ACTN4-modest lamellipodia and ACTN4-enriched lamellipodia, the moving speed of ACTN4-enriched lamellipodia (average speed of 0.37 ± 0.15 µm/min, n = 15) was significantly faster than that of ACTN4-modest lamellipodia (average speed of 0.07 ± 0.04 µm/min, n = 15). \*\**P*<0.01.

It is also notable that ACTN4-enriched lamellipodia have wavy structures at their leading edges. These wavy structures were recognized as phase-dark structures, similar to membrane ruffles, by phase-contrast microscopy. However, compared with canonical membrane ruffles observed in macrophage colony-stimulating factor (m-CSF)-stimulated macrophages and epidermal growth factor (EGF)-stimulated A431 cells, ruffle-like structures of ACTN4-enriched lamellipodia in BSC-1 cells appeared to be smaller in size and tightly stacked within the region of leading edges (Video 4). It has previously been shown that membrane ruffling in macrophages and A431 cells is resistant to PI3K inhibition, although the process of membrane ruffle closure to macropinosomes is PI3K-dependent [21,45]. Therefore, the ruffle-like structures of the ACTN4-enriched lamellipodial leading edges may be distinct in nature from the canonical membrane ruffles.

The long-term movie over several hours showed that BSC-1 cells expressing EGFP-ACTN4 dynamically migrate upon extending lamellipodia with phase-dark wavy leading edges. However, when lamellipodia were static or retracting, ruffle-like wavy structures were absent from the leading edges (Video 5). Thus, ruffle-like wavy structures of the lamellipodial leading edges may be hallmarks of progressive lamellipodia.

### CLEM revealed that ACTN4-enriched lamellipodia have multilayered membrane folds at their leading edges

To clarify the ultrastructural features of phase-dark wavy-edge lamellipodia observed under phase-contrast microscopy, we performed CLEM on BSC-1 cells expressing EGFP-ACTN4. First, BSC-1 cells transfected with pEGFP-ACTN4 were cultured on gridded glass coverslips in 35 mm dishes and fixed with 2% glutaraldehyde in a buffer. After washing with PBS, the specimens were observed and photographed with a Zeiss LSM710 confocal microscope using a Plan-Apochromat 40× water-immersion objective lens to identify the precise location of cells expressing EGFP-ACTN4. We then processed the specimen coverslips for scanning EM and observed the same area under the scanning EM. CLEM showed that ACTN4-enriched lamellipodia have a characteristic ultrastructure; the leading edges of ACTN4-enriched lamellipodia have multilayered membrane folds (Fig.5). Consistent with the phase-contrast microscopy results, membrane folds at the lamellipodial leading edges, where ACTN4 was particularly concentrated, were shorter but more densely present than membrane ruffles in RAW macrophages and A431 cells (Supplementary Fig. S5).

Consequently, it can be asserted that multilayered membrane folds at the lamellipodial leading edges are strictly distinct from the simple membrane ruffles observed in other cell types, although they can be broadly considered small ruffles. Thus, we found that ACTN4-enriched lamellipodia in BSC-1 cells possess a unique morphological feature characterized by multilayered ruffles at the leading edge. Based on this characteristic, we hereafter refer to them as “ruffle-edge lamellipodia” to distinguish them from conventional flat lamellipodia.

**Figure 5.**
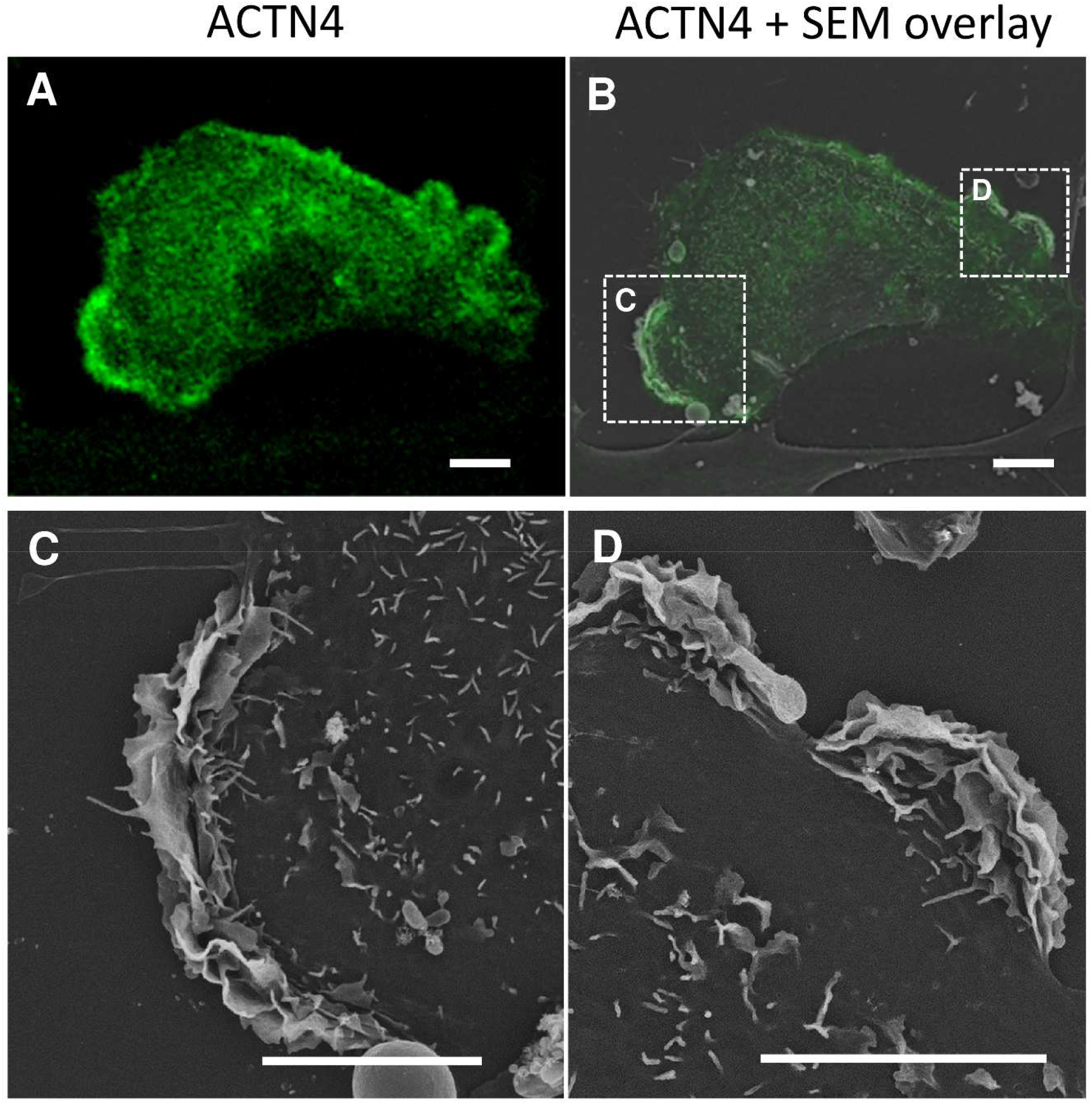
Correlative light-electron microscopy (CLEM) of EGFP-ACTN4-enriched lamellipodia. (A) Fluorescence microscopic image of EGFP-ACTN4 in a BSC-1 cell. (B) The ACTN4 image was overlaid on the scanning EM image of the same cell to identify ACTN4-enriched lamellipodia. Scanning electron micrographs of the boxed regions in B are magnified in the lower panels (C, D). Note that ACTN4-enriched lamellipodia have multilayered membrane folds at the leading edge. Scale bars, 10 μm.

### Formation of ACTN4-enriched ruffle-edges may require functional Rac1 switching

It is well-known that Rac1 activation is required for canonical lamellipodial formation [9,46,47]. However, the involvement of Rac1 activity in the formation of ACTN4-enriched ruffle-edge lamellipodia remains unknown. Therefore, we examined the impact of Rac1 mutant overexpression on ACTN4-enriched lamellipodia. BSC-1 cells were transfected with pmCitrine-fused wild-type Rac1 (Rac1 WT), a constitutively active GTP-lock mutant (Rac1 Q61L), or a dominant-negative GDP-lock mutant (Rac1 T17N). Following fixation, ACTN4 was detected by immunostaining with an anti-ACTN4 antibody, followed by incubation with an Alexa Fluor 594-labeled secondary antibody. In cells expressing Rac1 WT, ACTN4-enriched lamellipodia were frequently observed (Fig. 6A, left column). In contrast, constitutively active Rac1 Q61L expressing cells exhibited ACTN4-modest flat leading-edge lamellipodia along the outer perimeter (Fig. 6A, middle column). In Rac1 T17N expressing cells, both types of lamellipodia were rarely observed (Fig. 6A, right column). A quantitative assay counting cells with ACTN4-enriched lamellipodia in Rac1 WT-, Rac1 Q61L-, or Rac1T17N-expressing cells showed that both GTP-locked and GDP-locked Rac1 mutants significantly inhibited ruffle-edge lamellipodia formation compared with Rac1WT-expressing cells or non-expressing cells (Fig. 6B). To confirm the involvement of Rac1 activity in the formation of ruffle-edge lamellipodia, we applied optogenetic control of photoactivatable (PA)-Rac1 in live BSC-1 cells co-expressing EGFP-Rac1 and mCherry-PA-Rac1. Without photoactivation in the dark (Fig. 6C, PA-Rac1 OFF), cells expressing PA-Rac1 had ACTN4-enriched ruffle-edge lamellipodia, probably owing to endogenous wild-type Rac1 activity. Continuous photoactivation of PA-Rac1 by blue light illumination for several minutes changed the lamellipodia phenotype from ACTN4-enriched ruffle-edge lamellipodia to ACTN4-modest flat-edge lamellipodia (Fig. 6C, Rac1 ON 6 min, 22 min, Video 6). These results indicate that functional Rac1, capable of switching between GTP-bound active and GDP-bound inactive states, may be indispensable for ruffle-edge lamellipodia, although only the GTP-bound active state of Rac1 may be required for the extension of canonical flat lamellipodia.

**Figure 6.**
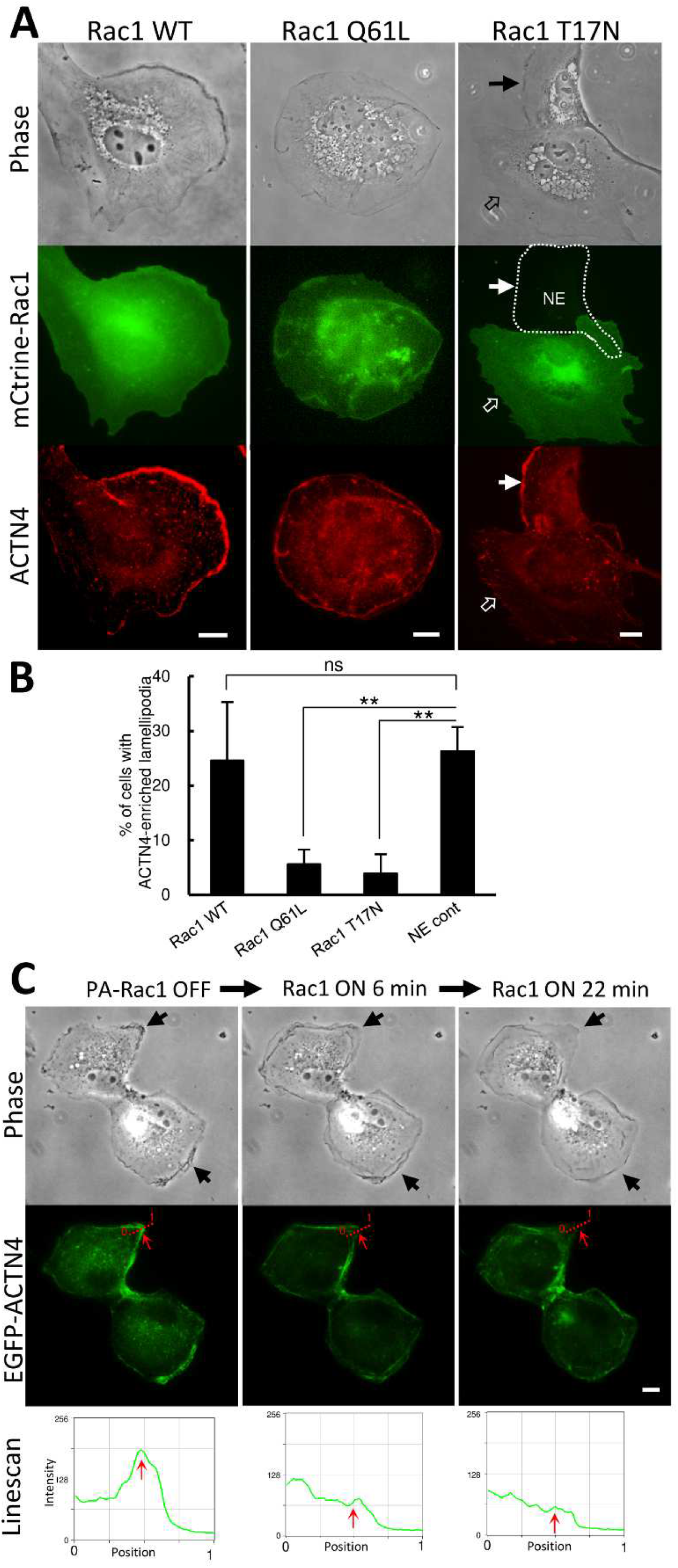
Rac1, which is capable of switching between GDP-bound and GTP-bound states, is indispensable for ACTN4-enriched lamellipodia. (A) BSC-1 cells were transfected with pmCitrine-Rac1 of wild type (WT), constitutively active mutant (Q61L), or dominant negative mutant (T17N). After fixation, ACTN4 was detected by immunostaining with an anti-ACTN4 antibody, followed by an Alexa Fluor 594-labeled secondary antibody. ACTN4-enriched lamellipodia are remarkable in cells expressing Rac1 WT. In cells expressing constitutively active Rac1 Q61L, flat leading-edge lamellipodia were prominent around the cell perimeter. Rac1 T17N expressing cells scarcely have lamellipodia (open arrow), although the next non-expressing cells have ACTN4-enriched lamellipodia (arrow). (B) Quantitation of the percentage of cells having ACTN4-enriched lamellipodia among Rac1 WT-, Rac1 Q61L-, Rac1 T17N-expressing BSC-1 cells, or non-expressing control (NE cont) cells (n=6 coverslips from two independent experiments. Expression of Rac1mutants (Q61L or T17N) significantly reduced the proportion of cells with ACTN4-enriched lamellipodia compared to non-expressing control cells (*P*<0.01, n=6), whereas Rac1 WT expression did not. (C) Representative time-lapse images showing that continuous photoactivation of PA-Rac1 changes the lamellipodia phenotype from ACTN4-enriched ruffle-edge lamellipodia to flat-edge lamellipodia (arrows in Phase). Linescan analysis shows the change in EGFP-ACTN4 fluorescence intensity at the red dotted line across the lamellipodia in the middle panels. Continuous activation of PA-Rac1 decreased EGFP-ACTN4 fluorescence intensity in lamellipodia. Red arrows in the middle images and linescan graphs indicate the same positions. Scale bars, 10 μm.

### The ruffle-edge lamellipodia are frequently observed in highly invasive cancer cells

We investigated whether the presence of ruffle-edge lamellipodia is cell type-specific using several representative cell lines: HeLa, MDCK-II, B16, A549, and MDA-MB-231 cells. In HeLa cells, lamellipodia were rarely observed. ACTN4 was predominantly localized to stress fibers and focal adhesions (Supplementary Fig. S6, upper panel). Although MDCK-II cells occasionally exhibited lamellipodia, their leading edges were not enriched with ACTN4 (Supplementary Fig. S6, middle panel). B16 mouse melanoma cells frequently exhibit well-developed canonical-type flat lamellipodia. The localization of ACTN4 to the leading edge was modest (Supplementary Fig. S6, lower panel). In human invasive lung cancer A549 cells, ACTN4-enriched lamellipodia were occasionally observed. We found that ACTN4-enriched lamellipodia significantly increased after stimulation with 100 nM phorbol 12-myristate 13-acetate (PMA) (Supplementary Fig. S7A). In live A549 cells expressing EGFP-ACTN4, we confirmed that ACTN4-enriched lamellipodia in PMA-stimulated A549 cells were perturbed by PI3K inhibition (Supplementary Fig.S7B). By scanning EM, we confirmed the formation of multilayered ruffles at the lamellipodial leading edges in PMA-stimulated A549 cells (Supplementary Fig.S7C).

To clarify the contribution of ACTN4-enriched ruffle-edge lamellipodia to cell migration, we observed the cell migration process during wound healing of a confluent monolayer culture scraped with a 200 µL pipette tip. Four hours after creating a wound on the coverslip monolayer culture of A549 cells, the cells were fixed with 4% PFA, immunostained for endogenous ACTN4, and observed by fluorescence microscopy and CLEM. Immunofluorescence analysis of ACTN4 showed that ACTN4 was enriched in the leading edge of lamellipodia, extending toward the wounded space. However, in the presence of LY294002, ACTN4-enriched lamellipodia were rarely observed (Supplementary Fig. S8A). Furthermore, CLEM of ACTN4 immunofluorescence and scanning EM clearly demonstrated that ACTN4-enriched lamellipodia have multilayered membrane folds at their leading edges (Supplementary Fig. S8B). Similarly, another highly invasive cancer cell line, human breast cancer MDA-MB-231, also showed ACTN4-enriched lamellipodia formation during wound healing in a PI3K-dependent manner (Supplementary Fig. S9). These findings suggest that ruffle-edge lamellipodia are not present in all cell types, but may be a characteristic feature of certain invasive cancer cells.

### The ruffle-edge lamellipodia formation and cell migration are suppressed by ACTN4 knockdown

Next, we examined the effect of ACTN4 knockdown on the formation of ruffle-edge lamellipodia and cell migration in A549 cells. Isoform-specific knockdown of ACTNs by ACTN4 or ACTN1 siRNAs was verified by immunofluorescence microscopy and immunoblotting (Supplementary Fig. S10). Five hours after scratching the A549 cell monolayer, cells extending ACTN4-enriched lamellipodia toward the wound space were observed in control siRNA-or ACTN1 siRNA-transfected cells, whereas cells treated with ACTN4 siRNAs did not show marked cell extension (Fig. 7A, Supplementary Fig. S10A). CLEM clearly demonstrated that ACTN4 knockdown cells showed only flat protrusions facing the wound space. However, ACTN1 siRNA did not show such an effect on ruffle-edge lamellipodia (Fig. 7B). Furthermore, the wound healing assay demonstrated that cell migration to the scratch wound was significantly reduced by ACTN4 knockdown (*P*<0.01, n=3), but not by ACTN1 knockdown (Fig. 7C). These results indicate a critical role for ACTN4 in cell migration through the formation of ruffle-edge lamellipodia.

**Figure 7.**
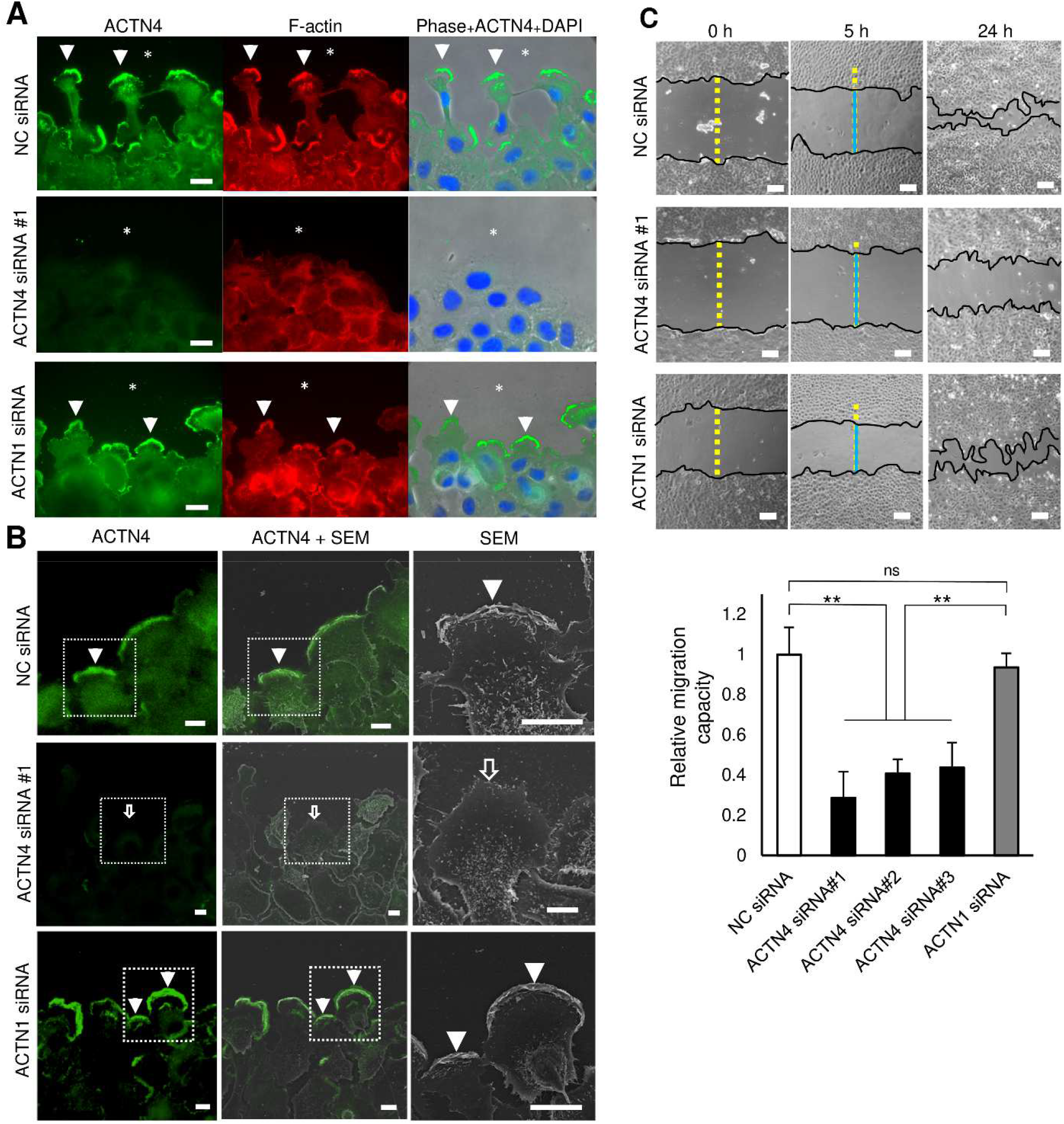
ACTN4 knockdown suppressed ruffle-edge lamellipodia formation and cell migration in A*549 cells. (A) A54*9 cells subconfluentl*y cultured on coverslips were transfected with negative control (NC) siRNA, ACTN4 siRNA (#1-3), or ACTN1 siRNA. After 48 h of transfection, the monolayer culture was scratched with a 200 µL pipette tip to make a wound and incubated for 5 h in the presence of 50 nM PMA. After fixation, ACTN4 was detected by immunofluorescence. F-actin and nuclei were stained with Alexa Fluor 568-phalloidin and DAPI, respectively. Fluorescence microscopy demonstrated that ACTN4-enriched lamellipodia extended to the scratched wound space of the monolayer culture in which cells were treated with NC siRNA or ACTN1 siRNA (arrowheads). However, lamellipodial extension was hardly observed in ACTN4 knockdown cells by ACTN4 siRNA#1. Asterisks denote the wounded space. Scale bars, 10 μm. (B) CLEM of ACTN4 fluorescence and scanning electron microscopy (SEM) reveals that isoform-specific knockdown of ACTN4 by siRNA inhibits ruffle-edge lamellipodia formation during cell migration. After fluorescence microscopy, cells were processed to scanning EM. The same area as ACTN4 fluorescence micrograph was photographed by SEM, and the two images were superimposed using Adobe Photoshop. Arrowheads indicate ACTN4-enriched ruffle-edge lamellipodia in A549 cells transfected with NC or ACTN1 siRNA. In ACTN4 KD cells by ACTN4 siRNAs, only flat thin lamellipodia were seen (open arrows). Similar results were obtained in cells transfected with ACTN4 siRNAs #2 and #3. Scale bars: 10 μm. (C) Wound healing assay demonstrating that ACTN4 knockdown perturbed the migration capacity of A549 cells. Representative images of scratched and recovering wounded areas at the indicated times (black lines) on confluent monolayers of cancer cells treated with NC siRNA, ACTN4 siRNA#1, or ACTN1 siRNA. Cell migration capacity was determined by measuring the migration distance by subtracting the width of the wounds at 5 h (blue lines) after making the wounds on cell monolayers from that at 0 h (yellow broken lines). Graph data are presented as fold-changes relative to the migration distance of NC siRNA-treated cells. ACTN4-KD cells showed a significantly lower migration capacity than ACTN1-KD or control cells (\**P*<0.05, n=3). Values are expressed as means ± SEM of three independent experiments.

### Possible role of the multilayered membrane ruffles at the lamellipodial leading edge in extracellular matrix degradation

Finally, we explored the functional significance of the characteristic layered membrane ruffles at the leading edge of lamellipodia. Certain invasive cancer cells invade by forming rod-like protrusions called invadopodia on the ventral surface of the cell, which degrade the ECM, including the basement membrane, by matrix metalloproteases (MPPs). If ruffle-edge lamellipodia are involved in cell invasion, it is possible that MMPs are also localized in the membrane ruffles at the leading edges. Therefore, we examined the localization of membrane type-1 matrix metalloproteinase (MT1-MMP, MMP14), a major physiological collagenase, in PMA-stimulated A549 cells by immunofluorescence microscopy. As expected, MT1-MMP and ACTN4 were abundantly colocalized in multilayered membranes at the lamellipodial leading edge (Fig. 8A, upper panel). Localization of MT1-MMP in the leading edge of lamellipodia was markedly diminished together with ACTN4 in A549 cells after LY294002 treatment (Fig.8A, lower panel).

**Figure 8.**
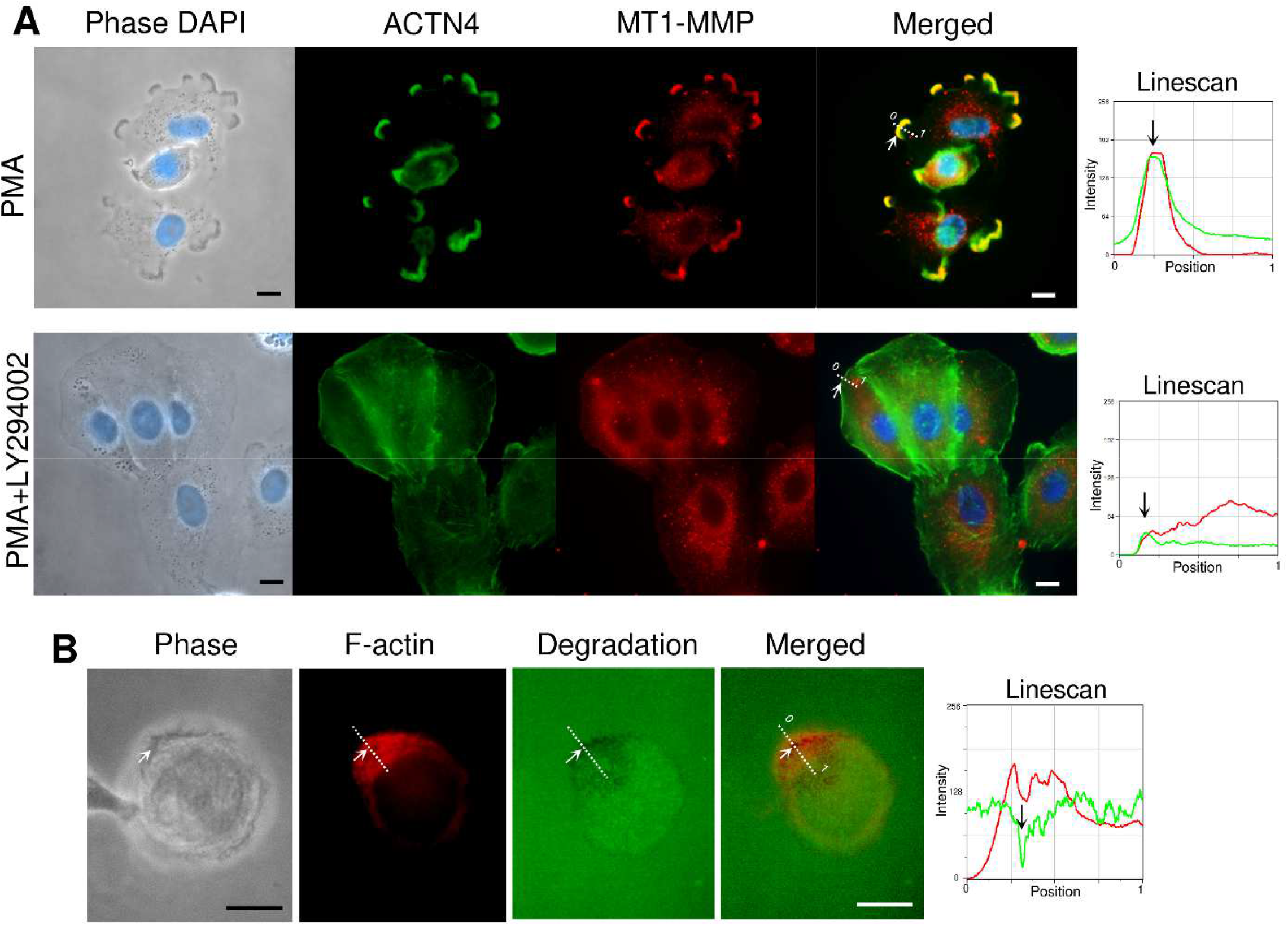
MT1-MMP localization and gelatin degradation activity of ruffle-edge lamellipodia. (A) PMA-stimulated A549 cells were immunolabeled for endogenous ACTN4 (green) and MT1-MMP (red). MT1-MMP localizes to ACTN4-enriched ruffle-edge lamellipodia (upper panel). Colocalization of ACTN4 and MT1-MMP is evident by the linescan analysis at the dotted line in the merged images. Arrows indicate the same position in the merged image and the linescan. After treatment with LY294002, both ACTN4 and MT1-MMP signals were markedly diminished (lower panel). DAPI, nuclear stain. (B) A549 cells were plated onto FITC-gelatin-coated coverslips and incubated for 3 h in the presence of 100 nM PMA. After fixation with 4% PFA, F-actin was stained with Alexa Fluor 568-phalloidin. FITC-gelatin degradation activity was observed as dark areas beneath lamellipodia tips. Scale bars, 10 μm.

Additionally, we examined whether A549 cells with ruffle-edge lamellipodia show focal ECM degradation activity on fluorescein isothiocyanate (FITC)-gelatin–coated coverslips. Three hours after plating A549 cells on FITC-gelatin coverslips, the cells produced gelatin degradation areas predominantly located at the leading edge of the lamellipodia (Fig. 8B). Taken together, these observations suggest that ruffle-edge lamellipodia may have ECM degradation activity by MT1-MMP, which localizes in multilayered membrane ruffles.

## Discussion

Actinin-1 (ACTN1) and actinin-4 (ACTN4), ubiquitously expressed in non-muscle cells, exhibit high amino acid similarity (87% identity) but show distinct localization and functional roles in cells [22,26,48]. Since its identification by Honda et al. (1998)[26], numerous studies have established that ACTN4 expression is closely associated with cancer invasion and metastasis [32,48–50]. However, the underlying molecular mechanisms are not fully understood.

In this study, we showed that subcellular localization of ACTN4 was most notable in dynamic lamellipodia, whereas ACTN1 was enriched in stress fibers and focal adhesions, and less frequently observed in lamellipodia. Moreover, ACTN4 overexpression significantly increased the proportion of cells with lamellipodia and, in turn, reduced the proportion of cells with stress fibers and focal adhesions. In contrast, overexpression of ACTN1 reduced the proportion of cells with lamellipodia. These results seem to be consistent with those of a previous study showing that ACTN4 overexpression enhances colorectal cancer cell invasion by suppressing focal adhesions and stress fibers [32]. Their study demonstrated that the recruitment of zyxin, a scaffolding protein for focal adhesions, is impaired by ACTN4 overexpression. The reduction in focal adhesions in ACTN4-overexpressing cells can be explained by zyxin-binding-deficient ACTN4 displacement with ACTN1 in focal adhesions.

Furthermore, our live-cell imaging of EGFP-ACTN4 expressing BSC-1 cells and kymograph analysis demonstrated that ACTN4-enriched lamellipodia showed more dynamic advancement than ACTN4-modest lamellipodia. Therefore, ACTN4 may exert its cell migration ability through the formation of ACTN4-enriched lamellipodia. Several studies have suggested a positive correlation between ACTN4 levels and cancer cell motility/invasion [31,51,52]. However, a direct association between ACTN4-enriched lamellipodia and cell migration has not yet been reported. Because phase-contrast microscopy showed that ACTN4-enriched lamellipodia had phase-dark wavy structures at their leading edges, we examined their ultrastructure using CLEM. As expected, CLEM demonstrated that ACTN4-enriched lamellipodia had a unique morphology of multilayered membrane folds at the leading edge. The membrane folds were smaller and more tightly stacked than the membrane ruffles observed in other cell types, including macrophages and EGF-stimulated A431 cells [22,53](Video 4, Fig. S5). Although conventional membrane ruffles raised by the upward vending of lamellipodia move to the dorsal surface of the cell and sometimes form circular ruffles, which are precursors to macropinosomes, the presence of layered membrane folds is restricted to a narrow area of the lamellipodial edges (Video 4, 5). More importantly, membrane ruffles in A431 cells and macrophages are resistance to PI3K-inhibitors [21,45], whereas the layered membrane folds of lamellipodial leading edges observed in this study are highly sensitive to PI3K inhibitors. Therefore, the morphology, function, and regulation of multilayered ruffles at the lamellipodial edges are likely distinct from those of conventional membrane ruffles. We considered ACTN4-enriched lamellipodia with multilayered ruffles at the leading edge as a new phenotype adapted to active cell migration and termed them “ruffle-edge lamellipodia” as one of the diverse cell membrane protrusions.

Our study demonstrated that ACTN4-enriched ruffle-edge lamellipodia are highly dependent on PI3K activity. PI3K inhibition by LY294002 drastically diminished ACTN4 in lamellipodia and prevented further extension of lamellipodia. Furthermore, high levels of PI(3,4,5)P_3_, a major product of class I PI3K, were observed in ruffle-edge lamellipodia using fluorescent protein-fused Akt-PH and Btk-PH domains. In contrast, PI3K inhibition did not perturb ACTN4 localization in stress fibers and focal adhesions, where no PI(3,4,5)P_3_ was detected. Phosphoinositides, particularly PI(4,5)P_2_ and PI(3,4,5)P_3,_ regulate the activity of several actin-binding proteins [16]. It has been reported that both PI(4,5)P_2_ and PI(3,4,5)P_3_ bind to the calponin homology 2 (CH2) domain of ACTN1 but differentially regulate actinin function by modulating the structure and flexibility of the molecule [43,44]. Interestingly, PI(3,4,5)P_3_ binding to ACTN1 inhibits actin-bunding activity and the interaction between ACTN1 and integrins at focal adhesions [42,54]. As actinin isoforms share highly homologous amino acid sequences, including the CH2 domain, the ACTN4 association with ruffle-edge lamellipodia may be mediated by the direct binding of PI(3,4,5)P_3_ to the CH2 domain of ACTN4. PI(3,4,5)P_3_ -bound ACTN4 may be able to respond to more dynamic movements of the actin network in ruffle-edge lamellipodia. It was also reported that class I PI3K binds to actinin through the p85 subunit [55]. Therefore, ACTN4 can be considered an important downstream effector of class I PI3K in the formation of ruffle-edge lamellipodia, although other PI3K downstream molecules may also be implicated in this process.

It is well-known that Rac1 activation plays a critical role in lamellipodial extension through branched actin network formation by activating the WAVE regulatory complex [17,46,47]. In BSC-1 cells expressing the constitutively active Rac1 Q61L mutant, flat lamellipodia were prominently observed, whereas ACTN4-enriched lamellipodia were rarely observed. In cells expressing a dominant-negative Rac1 mutant (Rac1T17N), neither flat nor ruffle-edge lamellipodia were observed. Only BSC-1 cells expressing wild-type Rac1 exhibited ruffle-edge lamellipodia formation. Consistently, the optogenetics of PA-Rac1 also revealed that ACTN4-enriched ruffle-edge lamellipodia changed to ACTN4-modest flat lamellipodia by continuous Rac1 photoactivation. We have previously shown that the formation of phagosomes and macropinosomes by cup-shaped lamellipodia is driven by Rac1 ON-OFF switching [37,39]. Similarly, spatiotemporally regulated ON-OFF switching of Rac1 may be indispensable for membrane fold formation at the leading edge of ACTN4-enriched lamellipodia, although only the Rac1 ON state may be sufficient for forming simple flat lamellipodia.

Importantly, we confirmed the presence of ruffle-edge lamellipodia in some invasive cancer cells, including A549 lung cancer and MD-MBA-231 human breast cancer cells, but not in all cell types. Because both A549 and MD-MBA-231 cells are known to be representative of highly invasive cancer cells, ACTN4-enriched ruffle-edge lamellipodia may contribute to their invasiveness. ACTN4 knockdown experiments using ACTN4 siRNAs demonstrated that ACTN4 knockdown greatly suppressed ruffle-edge lamellipodium formation and migration in A549 cells. Thus, it is likely that ACTN4 is essential for progressive cell migration by extending ruffle-edge lamellipodia.

Invasive cancer cells cultured on an artificial extracellular matrix have invadopodia, which are other structures involved in cell invasion. Invadopodia are rod-shaped actin-rich membrane protrusions that arise from the ventral side of the cell and degrade the basal matrix via matrix metalloproteinases (MMPs) [56,57].

Similar to lamellipodia formation, invadopodia are formed through the N-WASP (but not WAVE1 and 2)-Arp2/3 pathway and many actin-binding proteins [58]. Interestingly, it has been reported that overexpression of ACTN4, but not ACTN1, promotes invadopodia formation [59]. In addition, class I PI3K signaling has been shown to be involved in the formation of invadopodia in breast cancer cells [60]. Thus, invadopodia and ACTN-4-enriched ruffle-edge lamellipodia share similarities in the molecular mechanisms of their formation. However, based on their morphology, their functional roles are likely to be distinct. Our live-cell and CLEM imaging demonstrated that ACTN4-enriched lamellipodia have multilayered leading edges and that ACTN4-enriched ruffle-edge lamellipodia aggressively extend in the lateral direction. In contrast, invadopodia extending from ventral cell membranes degrade the basement membrane during cancer cell invasion. Invadopodia and ruffle-edge lamellipodia are likely different cancer invasion modalities with distinct roles: penetration of the basement membrane and invasion of the interstitial tissue by degrading extracellular matrix components, respectively. Considering the functional role of multilayered membrane ruffles at the leading edge of lamellipodia, the increase in membrane area due to the formation of multilayered folds may be advantageous for efficient ECM degradation by MT1-MMP to advance in the interstitium.

Lamellipodia have been recognized as two-dimensional flat cell protrusions; however, it has recently been reported that lamellipodia in zebrafish keratocytes may modulate the speed and direction of cell migration by causing three-dimensional morphological variations [61]. This study, together with ours, underscores that cell migration using lamellipodia extension also has various modes, depending on the cell type and surrounding environment. In a wide range of physiological, developmental, and disease-related processes, various modes of cell migration may be appropriately employed. There is also diversity in membrane ruffles among cell types, and at present, distinct types of membrane ruffles with different functions and structural characteristics seem to be conflated; therefore, they should be clearly classified. The discovery of multilayered ruffle-edge lamellipodia with ECM degradation activity challenges the conventional concepts of lamellipodia and membrane ruffles and will deepen our comprehensive understanding of the mechanisms of cancer invasion and metastasis. Prospectively, in vivo analysis of the three-dimensional construction and motility of multilayered ruffle-edge lamellipodia will hopefully lead to the elucidation of the mechanisms that enhance the invasive potential of cancer cells and to their suppression.

## Conclusions

In conclusion, ACTN4 plays an essential role in ruffle-edge lamellipodia formation in a PI3K-dependent manner. Given its highly characteristic traits, we consider ACTN4-enriched ruffle-edge lamellipodia as a new invasive mode of lamellipodia in certain cell types. Our study emphasizes that ACTN4 may play a critical role in the ability of cells to invade/migrate in the lateral direction through ruffle-edge lamellipodia formation. Further characterization of ACTN4-enriched ruffle-edge lamellipodia would greatly aid in the development of new criteria for cancer cell migration/invasion.

## Abbreviations

ACTN4: actinin-4
Arp2/3: Actin-related protein 2/3
EGFP: enhanced green fluorescent protein
CLEM: correlative light-electron microscopy
PH domain: pleckstrin homology domain
PI3K: phosphoinositide 3-kinase
PI(4,5)P_2_: phosphatidylinositol 4,5-bisphosphate
PI(3,4,5)P_3_: phosphatidylinositol 3,4,5-triphosphate
PMA: phorbol 12-myristate 13-acetate
N-WASP: neuronal Wiskott-Aldrich syndrome protein
WAVE: WASP family verprolin-homologous protein
MT1-MMP: membrane-type 1 matrix metalloproteinase.

## Acknowledgments

We are grateful to Drs. Joel A. Swanson (University of Michigan), Klaus Hahn (UNC-Chapel Hill), and Carol Otey (UNC-Chapel Hill) for providing plasmids. We also thank Dr. Eugene Park, Dr. Katsuya Miyake, Mr. Toshitaka Nakagawa, Mr. Kazuhiro Yokoi, and Ms. Yukiko Iwabu for their assistance and advice.

This study was supported by Grants in-Aid from the JSPS (#23K06306 to NA, #20K07245 to YE).

## CRediT authorship contribution statement

### Haruka Morishita

**Visualization, Validation**, Methodology, Investigation, Formal analysis, Data curation, Writing – original draft, Writing - review & editing. **Kawai Katsuhisa:** Methodology, Investigation, Data curation, Writing - review & editing. **Youhei Egami:** Methodology, Investigation, Data curation, Writing - review & editing, Funding acquisition. **Kazufumi Honda:** Resources, Validation, Writing - review & editing. **Nobukazu Araki:** Conceptualization, Project administration, Resources, Validation,Visualization, Investigation, Data curation, Writing - original draft; Writing - review & editing; Funding acquisition, Supervision.

### Declaration of Generative AI and AI-assisted technologies in the writing process

During the preparation of this work, the authors used Paperpal, an AI-assisted academic writing tool, in order to improve readability and language. After using this tool, the authors reviewed and edited the content as needed and take full responsibility for the content of the publication.

### Data availability

Most of the data can be found in this article and in the Supporting Information. Additional data are available upon request.

## Appendix A. Supplementary information

**Video 1. Live-cell movie showing the effect of LY294002 on EGFP-ACTN4 localization in BSC-1 cells**. Phase-contrast and fluorescence images of BSC-1 cells expressing EGFP-ACTN4 and mCherry-Lifeact were acquired at 15 s intervals for 20 min. EGFP-ACTN4 in the leading edges of lamellipodia markedly decreased within 5 min of the addition of LY294002 (white arrows), whereas that in the stress fibers and focal adhesions was sustained (blue arrows). The elapsed time after the addition of LY294002 is shown in the upper-left corner of the frame. 90× speed. Scale bars: 10 μm.

**Video 2. Live-cell movie showing the effect of LY294002 on lamellipodia enriched with PI(3**,**4**,**5)P**_**3**_ **monitored with YFP-Btk PH domain (right frame)**. PI(3,4,5)P_3_ is found to be enriched at the leading edge of the lamellipodia. LY294009 was added at the time point 8 min. The addition of LY294002 decreased PI(3,4,5)P_3_ levels in lamellipodia. The corresponding phase-contrast image is shown on the left-hand side of the figure. 90× speed. Scale bars: 10 μm.

**Video 3. Live-cell movie of EGFP-ACTN4 (right) and the corresponding DIC (left) images in BSC-1 cells**. The cell on the right side has an ACTN4-enriched lamellipodium, in which the leading edge shows a wavy appearance (arrow in the DIC images). The ACTN4-enriched lamellipodium (arrow) progressively extends, whereas the ACTN4-modest flat lamellipodia-like protrusions (open arrows) in the cell on the left side are not motile. 90× speed. Scale bars: 10 μm.

**Video 4. Phase-dark wavy structures at the lamellipodial leading edge in BSC-1 cells are distinct from membrane ruffling in m-CSF-stimulated RAW264 cells and EGF-stimulated A431 cells**. Membrane ruffles observed in RAW and A431 cells are mostly single lamellar structures originating from the cell periphery and move to the dorsal surface (black arrows). Wavy structures in BSC-1 cells are smaller and restricted to the narrow region of the lamellipodial leading edges (red arrows). 180× speed. Scale bars: 10 μm.

**Video 5. Long-term movie of an EGFP-ACTN4-expressing BSC-1 cell showing active cell migration through extending lamellipodia with phase-dark wavy edges**. After confirming EGFP-ACTN4 expression, phase-contrast images of the cell were acquired at 30 s intervals for 17 h. This ACTN4-expressing cell frequently forms lamellipodia, but their presence is temporary. The playback speed is displayed in the video frame. Scale bars: 10 μm.

**Video 6. Optogenetic activation of PA-Rac1 induces a morphological change from ACTN4-enriched ruffle-edge lamellipodia to ACTN4-modest flat lamellipodia exhibiting larger ruffling movement**. BSC-1 cells co-transfected with EGFP-ACTN4 and mCherry-LOV2-Rac1Q61L were subjected to PA-Rac1 photoactivation by blue light irradiation. Before photoactivation, the cells have ACTN4-enriched ruffle-edge lamellipodia (arrows), which is a phenotype of endogenous wild-type Rac1. Photoactivation of PA-Rac1 transformed ruffle-edge lamellipodia to flat lamellipodia within several minutes. The elapsed time after PA-Rac1 photoactivation is shown in the frame. 90× speed. Scale bars: 10 μm.

---

**Supplementary Figure S1.**
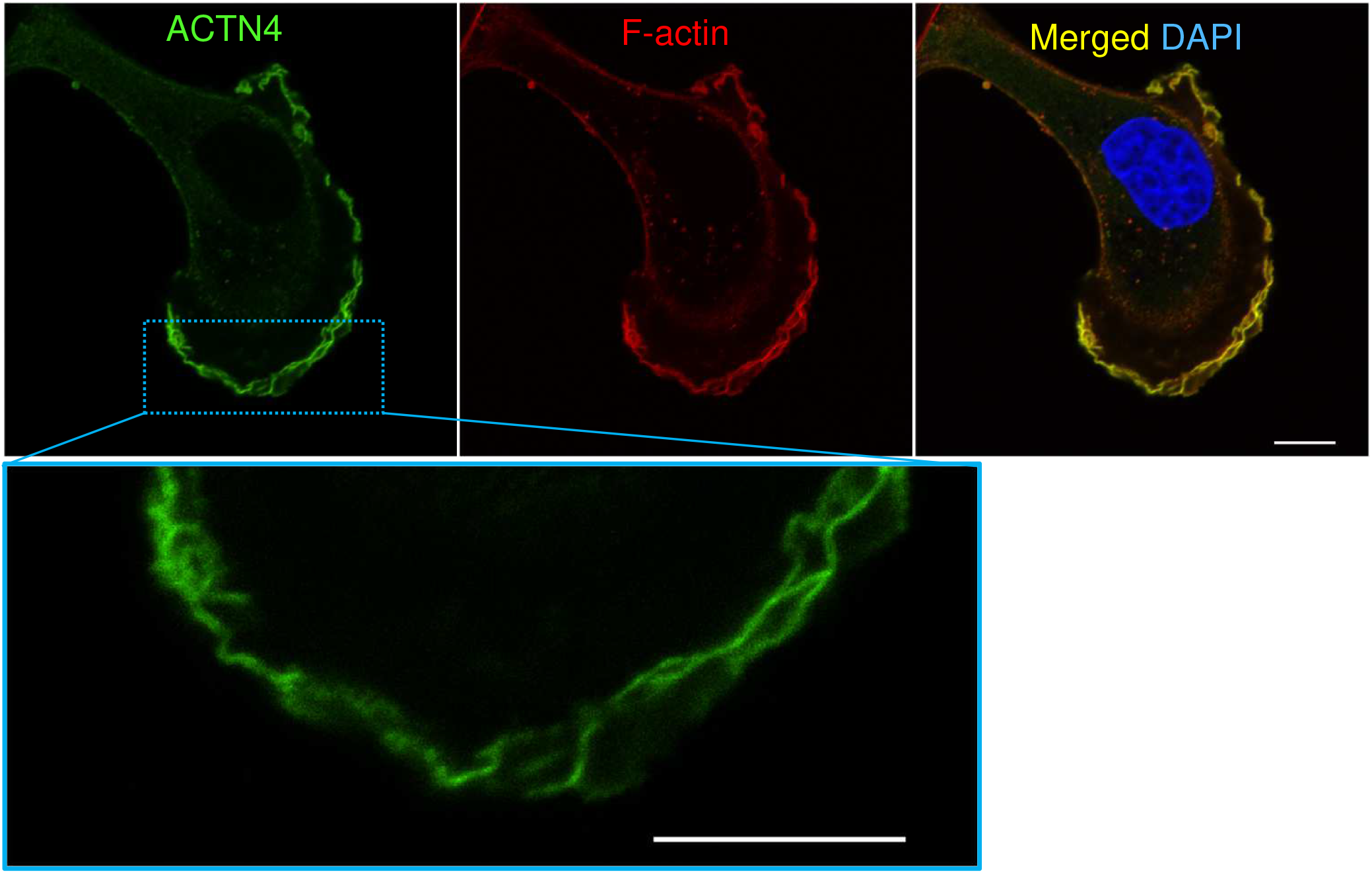
High-resolution confocal microscopy showing the representative localization of ACTN4 in L-type BSC-1 cells. BSC-1 cells transfected with EGFP-ACTN4 were fixed with 4% PFA. F-actin was stained with Alexa Fluor 568 phalloidin. Nuclei were stained with DAPI. High-resolution images were acquired with the Nikon AX NSPARC confocal laser microscope system. ACTN4 was observed to be associated with wavy membranous structures at the leading edge of lamellipodium. Actin stress fibers were rarely observed in L-type cells. The blue-boxed region was enlarged in the lower frame. Scale bars: 10 μm.

**Supplementary Figure S2.**
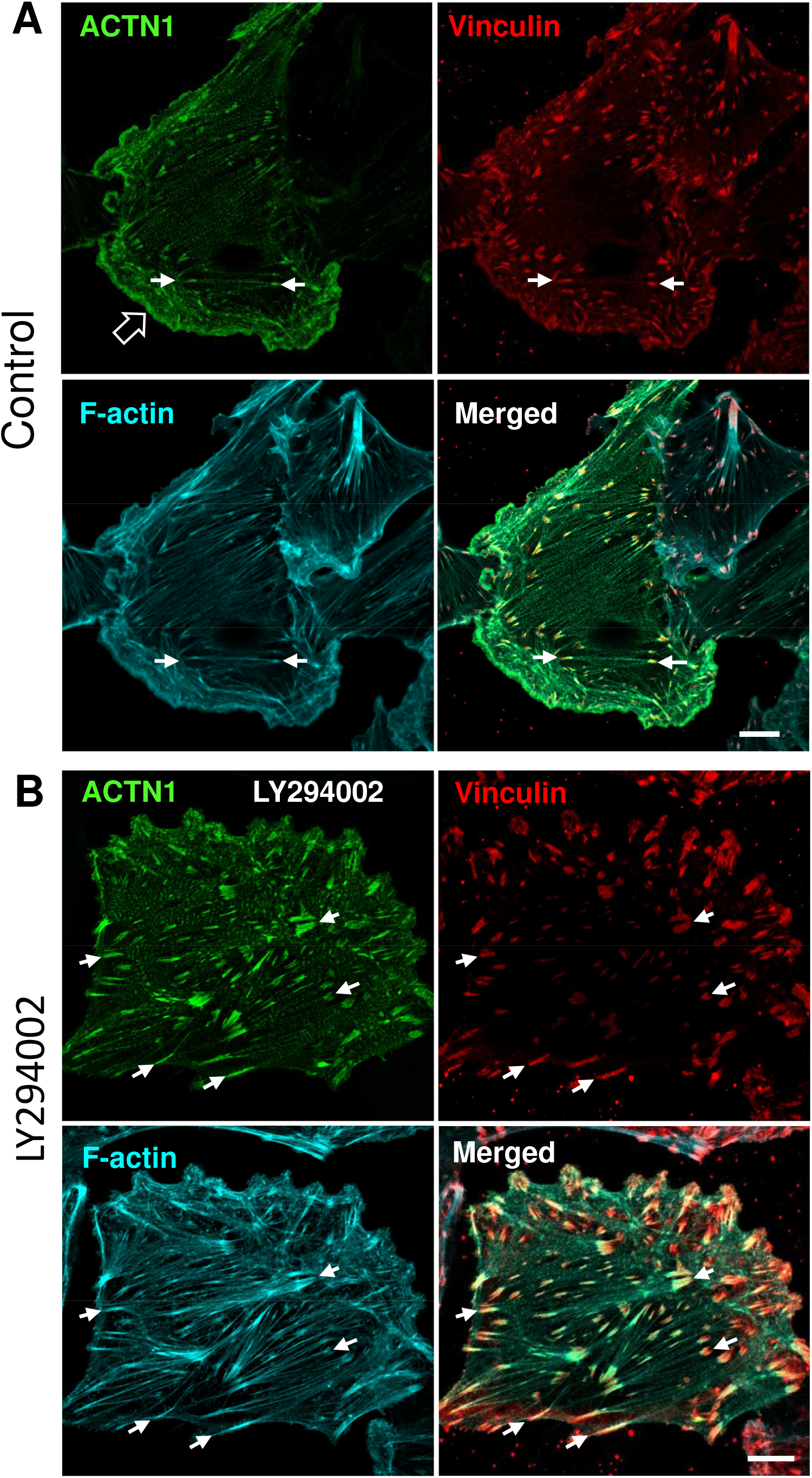
High-resolution confocal microscopy showing the representative localization of ACTN1 in control BSC-1 cells (A) and LY294002-treated cells (B). BSC-1 cells transfected with EGFP-ACTN1 were treated with 0.1% DMSO or 25 μM LY294002, followed by fixation with 4% PFA. Cells were immunostained with an anti-vinculin antibody, which labels focal adhesions. F-actin was stained with Acti-stain 670. Images were acquired with the Nikon AX NSPARC confocal laser microscope system. (**A**) ACTN1 was mainly localized to actin stress fibers and focal adhesions (arrows) in control BSC-1 cells. Localization of ACTN1 to lamellipodia (open arrow) was modest. (**B**) The localizations of ACTN1 in actin stress fibers and focal adhesions (arrows) was not diminished by the LY294002 treatment. Scale bars: 10 μm.

**Supplementary Figure S3.**
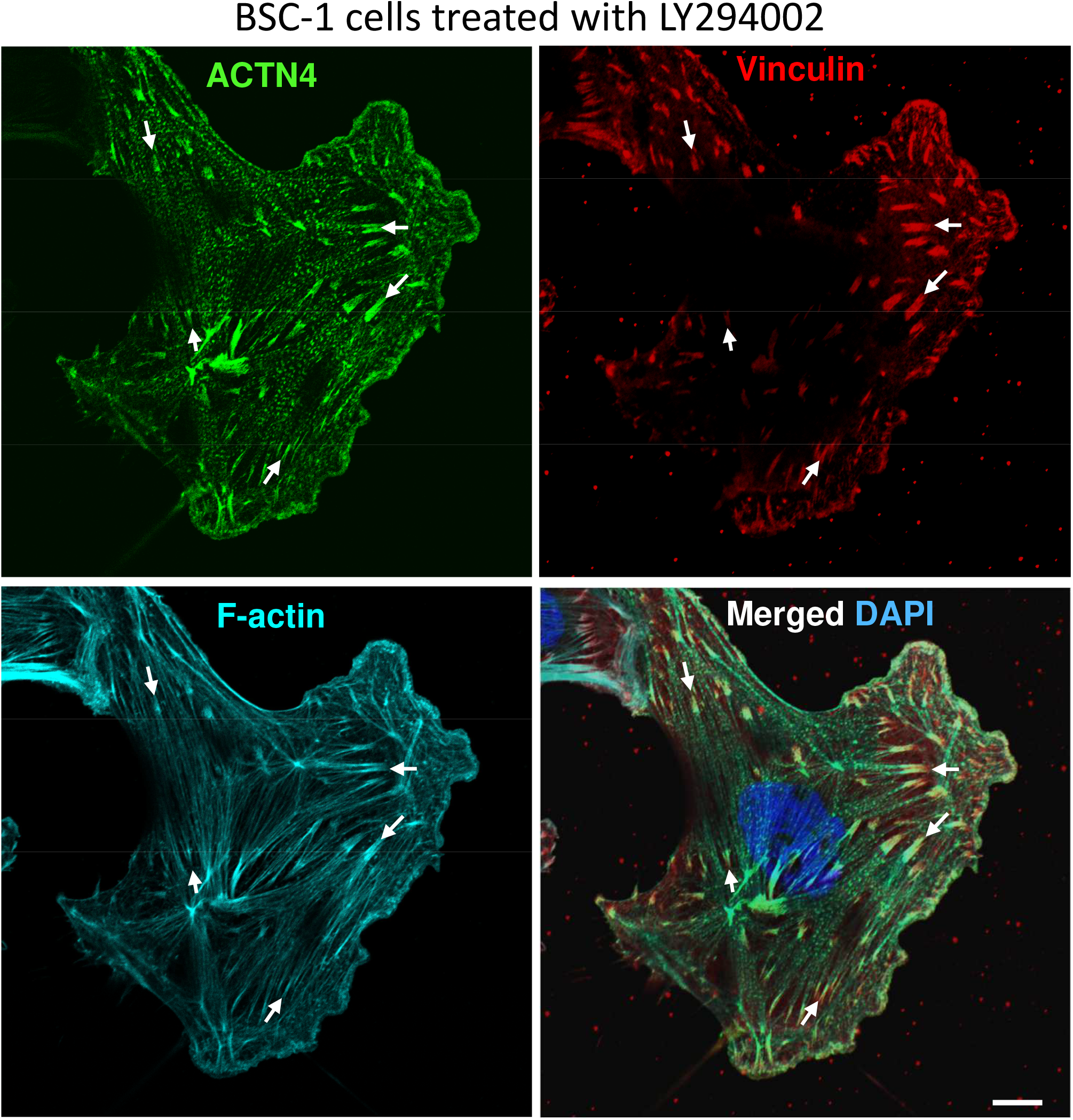
High-resolution confocal microscopy showing the representative localization of ACTN4 in LY294002-treated BSC-1 cells. BSC-1 cells transfected with EGFP-ACTN4 were treated with 25 µM LY294002 for 30 min and fixed with 4%PFA. Cells were immunostained with an anti-vinculin antibody, which labels focal adhesions (red). F-actin was stained with Acti-stain 670 (cyan). Nuclei were stained with DAPI (blue). Images were acquired with the Nikon AX NSPARC confocal laser microscope system. After LY294002 treatment, the localization of ACTN4 in lamellipodia was markedly diminished, while ACTN4 in actin stress fibers and focal adhesions (arrows) was sustained. Scale bars: 10 μm.

**Supplementary Figure S4.**
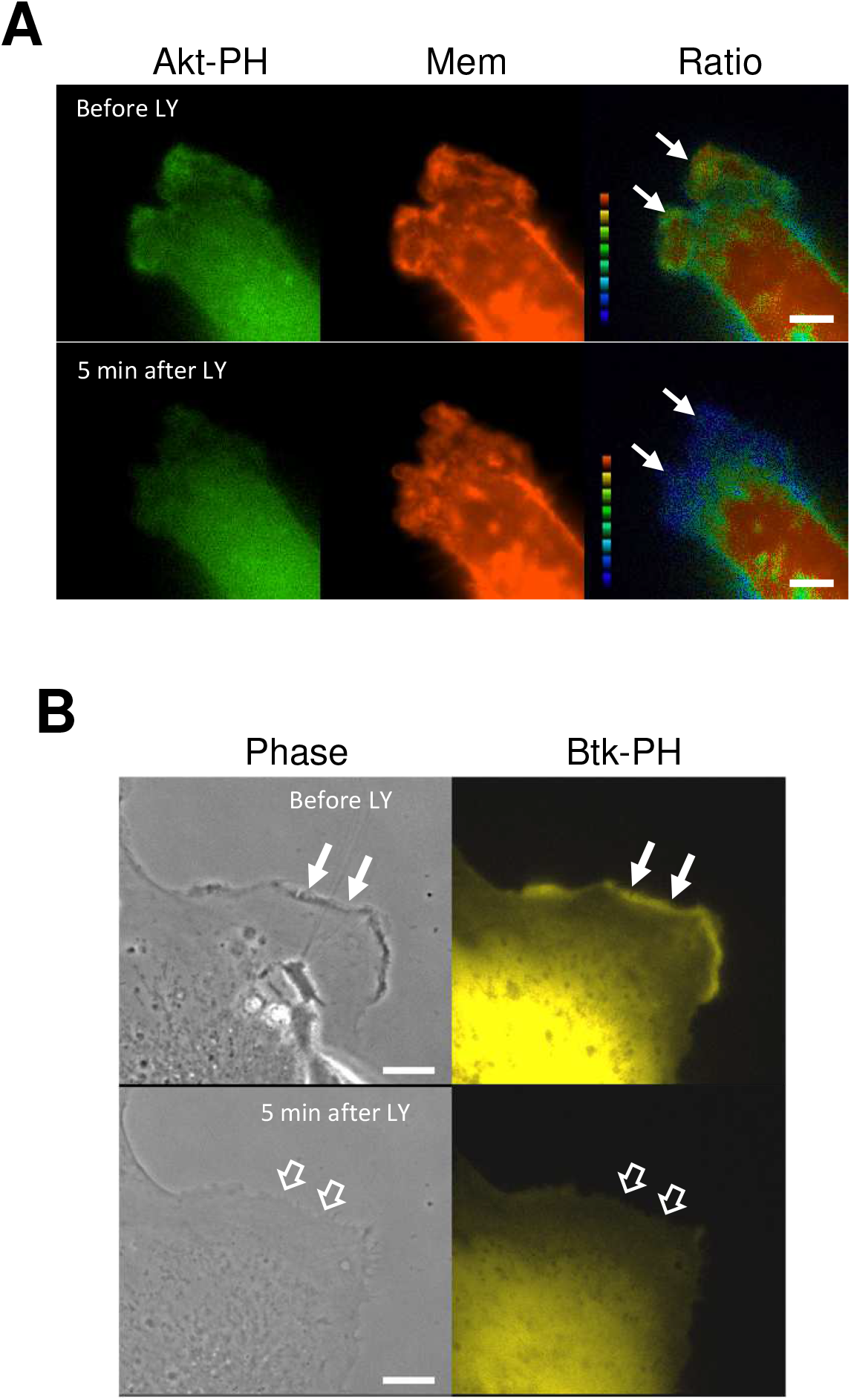
Effect of LY294002 on 3’ phosphoinositide in lamellipodial leading edges. (**A**) BSC-1 cells were co-transfected with mCitrine-Akt-PH and mCherry-membrane (Mem). Ratio images of Akt-PH/Mem showing that PI(3,4,5)P_3_/PI(3,4)P_2_ concentrations in the plasma membrane were created by the MetaMorph imaging system as previously described [21]. Note that PI(3,4,5)P_3_/PI(3,4)P_2_ concentrations in the membrane of lamellipodial leading edges (arrows) decreased by the addition of 25 μM LY294002 (LY). (**B**) Live-cell microscopy of a BSC1 cell showing the effect of LY294002 on lamellipodia enriched with PI(3,4,5)P_3_ monitored with YFP-Btk PH domain. In BSC-1 cells expressing YFP-Btk PH domain, PI(3,4,5)P_3_ was detected at the leading edge of lamellipodium (arrows). After the addition of LY, PI(3,4,5)P_3_ markedly diminished in the lamellipodium. The corresponding phase-contrast images are shown in the left frame. It is also notable that the phase-dark structures at the leading edge of the lamellipodium disappeared after the addition of LY (open arrows). Scale bars: 10 μm. A corresponding movie is available in the supplementary materials (**Video 2**).

**Supplementary Figure S5.**
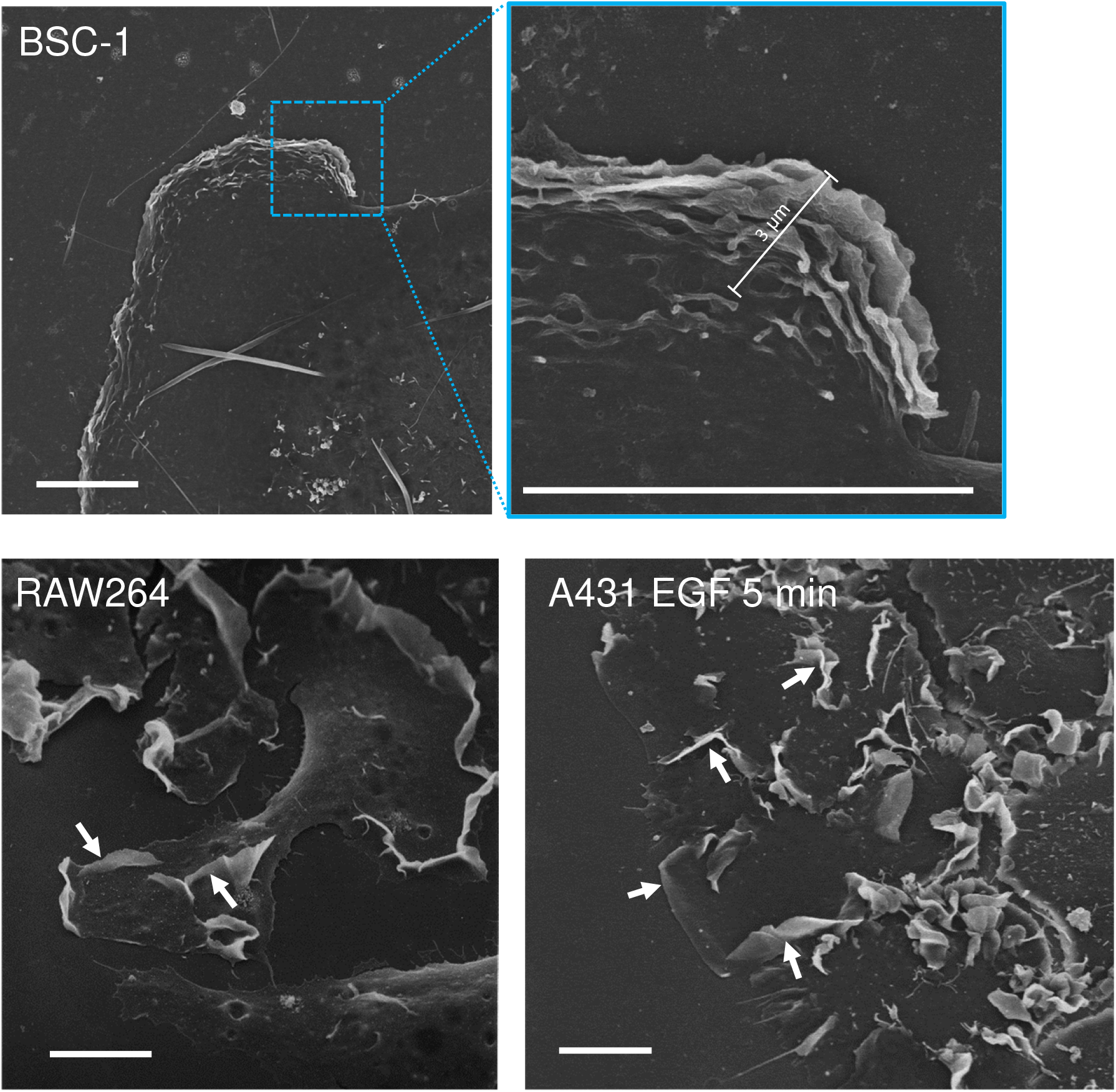
Scanning EM showing that ruffle-edge lamellipodia in BSC-1 cells are distinct from membrane ruffles arising from flat lamellipodia in RAW264 macrophages and EGF-stimulated A431 cells. Multilayered membrane ruffles of the lamellipodial leading edge in BSC-1 cells are smaller and more tightly stacked than membrane ruffles in RAW264 and EGF-stimulated A431 cells. Membrane ruffles arising from flat lamellipodia extend to the dorsal surface of RAW264 and A431 cells (arrows), whereas the presence of layered membrane ruffles is restricted to within ∼3 μm from the leading edge of the lamellipodium (upper right enlarged image of the boxed area) in BSC-1 cells. Scale bars: 10 μm.

**Supplementary Figure S6.**
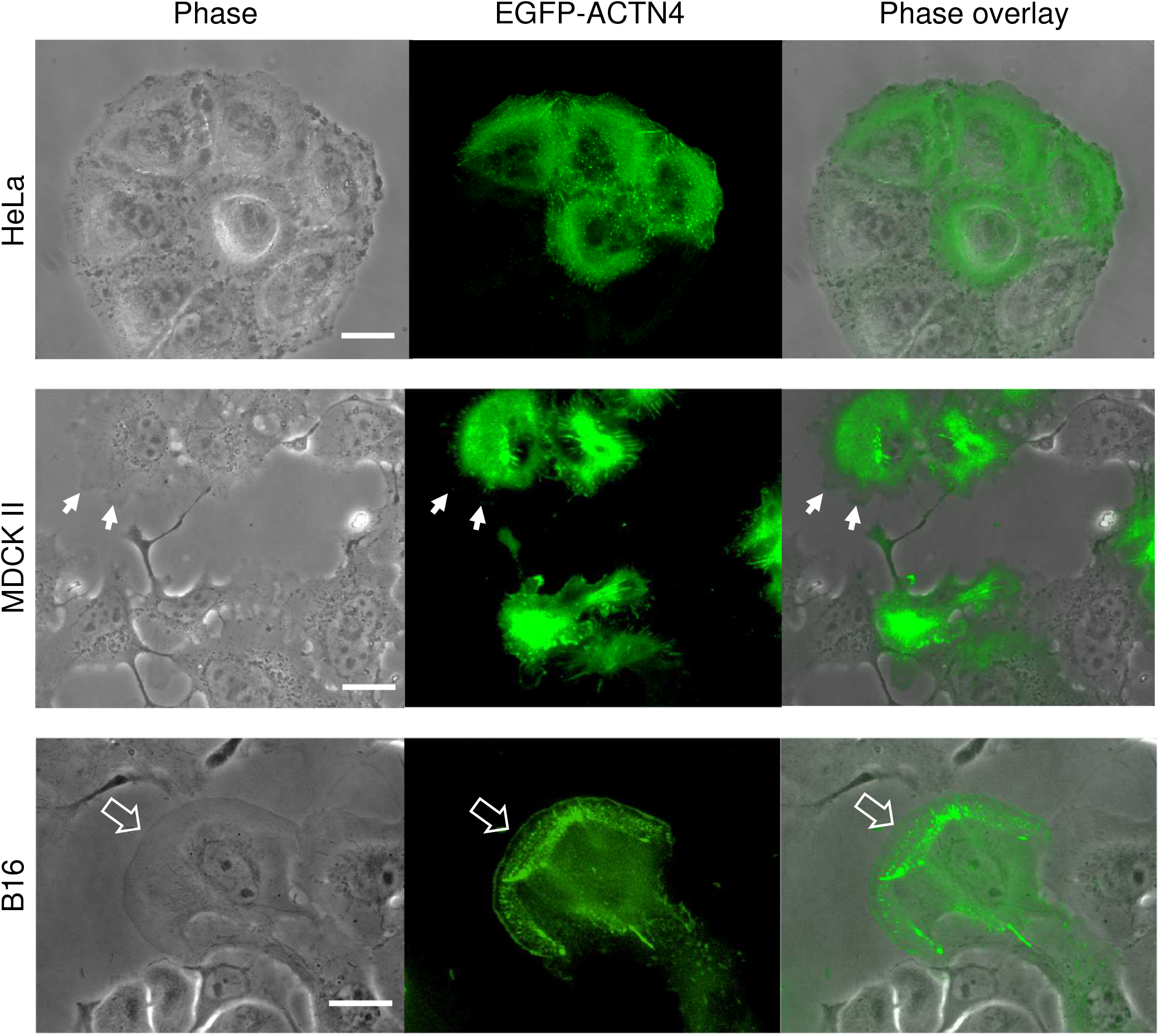
ACTN4-enriched lamellipodia were rarely observed in HeLa, MDCK II, or B16 cells. Live cells transfected with EGFP-ACTN4 were observed by phase-contrast and fluorescence microscopy. In HeLa cells, ACTN4 was predominantly localized in stress fibers and focal contacts. Although MDCK II cells frequently showed the extension of lamellipodia (arrows), they did not have ACTN4-enriched ruffle edge. B16 cells showed well-developed lamellipodia (open arrow), but the ACTN4 localization in their flat leading edges was modest. Scale bars: 10 µm.

**Supplementary Figure S7.**
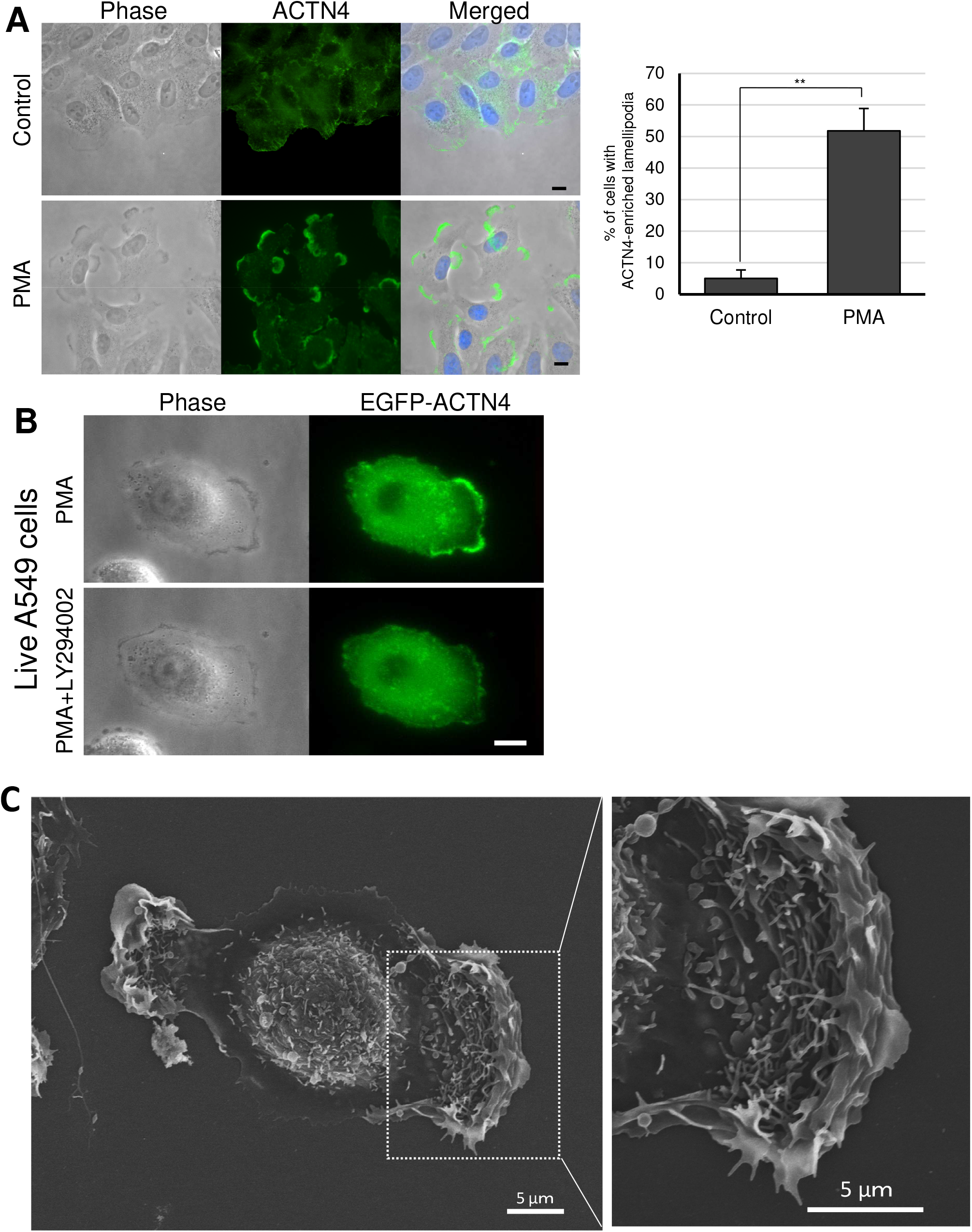
ACTN4-enriched ruffle-edge lamellipodia in A549 cells stimulated with PMA. (**A**) Immunofluorescence microscopy showing that ACTN4-enriched lamellipodia formation in A549 cells was markedly enhanced by 100 nM PMA treatment for 30 min. Quantitation by counting cells with ACTN4-enriched lamellipodia among >100 cells/coverslip (right graph). Percentages of A549 cells having ACTN4-enriched lamellipodia were significantly increased by the PMA-treatment. Each bar represents the mean ±SEM of 6 coverslips collected from two-independent experiments. **P<0.01. (**B**) Live-cell imaging of A549 cells transfected with EGFP-ACTN4 showing that ACTN4-enriched lamellipodia induced by PMA treatment (upper column) were diminished after the addition of LY294002 (lower column). Scale bar: 10 µm. (**C**) Scanning EM showing that lamellipodia have multilayered membrane folds at their leading edges in PMA-stimulated A549 cells. A magnified view of the boxed area is shown in the right frame. Scale bars: 5 µm.

**Supplementary Fig. S8.**
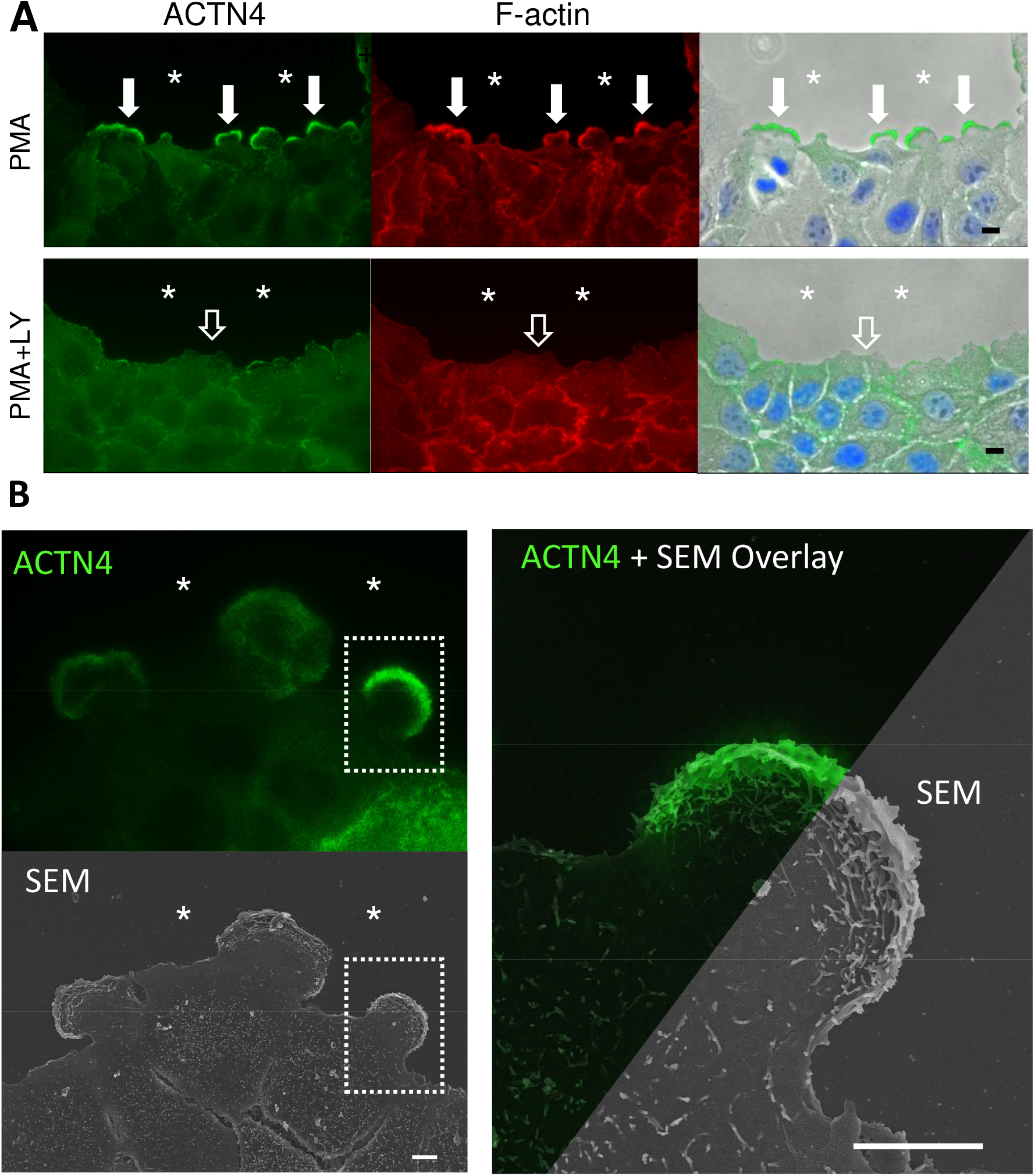
Formation of ACTN4-enriched ruffle-edge lamellipodia during wound healing in human non-small cell lung cancer A549 cell monolayer culture. (**A**) Three hours after creating a wound on the monolayer culture, the cells were fixed with PFA. Endogenous ACTN4 was immunostained with an anti-ACTN4 antibody, followed by Alexa Fluor 488-labeled secondary antibody (green). F-actin was stained with Alexa Fluor 568-phalloidin (red). Nuclei were stained with DAPI (blue). In PMA-stimulated A549 cells, ACTN4-enriched lamellipodia (arrows) were formed in migrating cells toward the wounded space (asterisks). The formation of ACTN4-enriched lamellipodia was inhibited by LY294002 (LY) treatment (open arrows). (**B**) CLEM reveals that ACTN4-enriched lamellipodia protruding toward the wounded space (asterisks) have multilayered membrane folds at their leading edges. A magnified view of the boxed area is shown in the right frame. Scale bars: 10 μm.

**Supplementary Figure S9.**
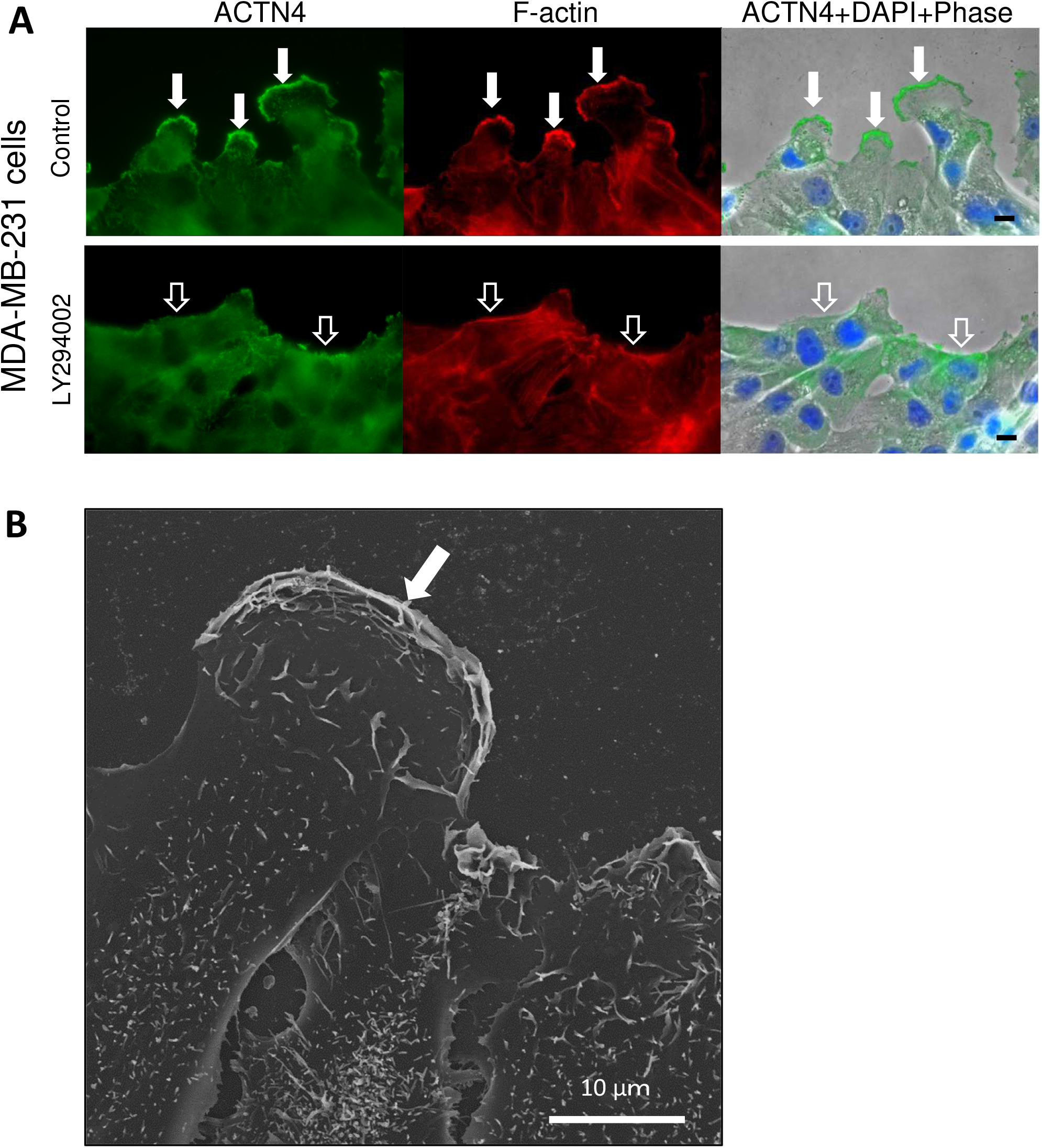
Formation of ACTN4-enriched ruffle-edge lamellipodia during wound healing of human invasive breast cancer MDA-MB-231 cell monolayer culture. (A) Three hours after creating a wound on the monolayer culture, cells were fixed and stained with anti-ACTN4 (green) and Alexa Fluor 564-phalloidine (red). In control MDA-MB-231 cells, ACTN4-enriced lamellipodia were formed in migrating cells toward the wounded space (arrows). ACTN4-enriched lamellipodia were not observed after LY294002 treatment (open arrows). Scale bars: 10 μm. (B) Scanning EM reveals that lamellipodia extending toward the wounded space have multilayered membrane folds (arrow). Scale bars: 10 μm.

**Supplementary Figure S10.**
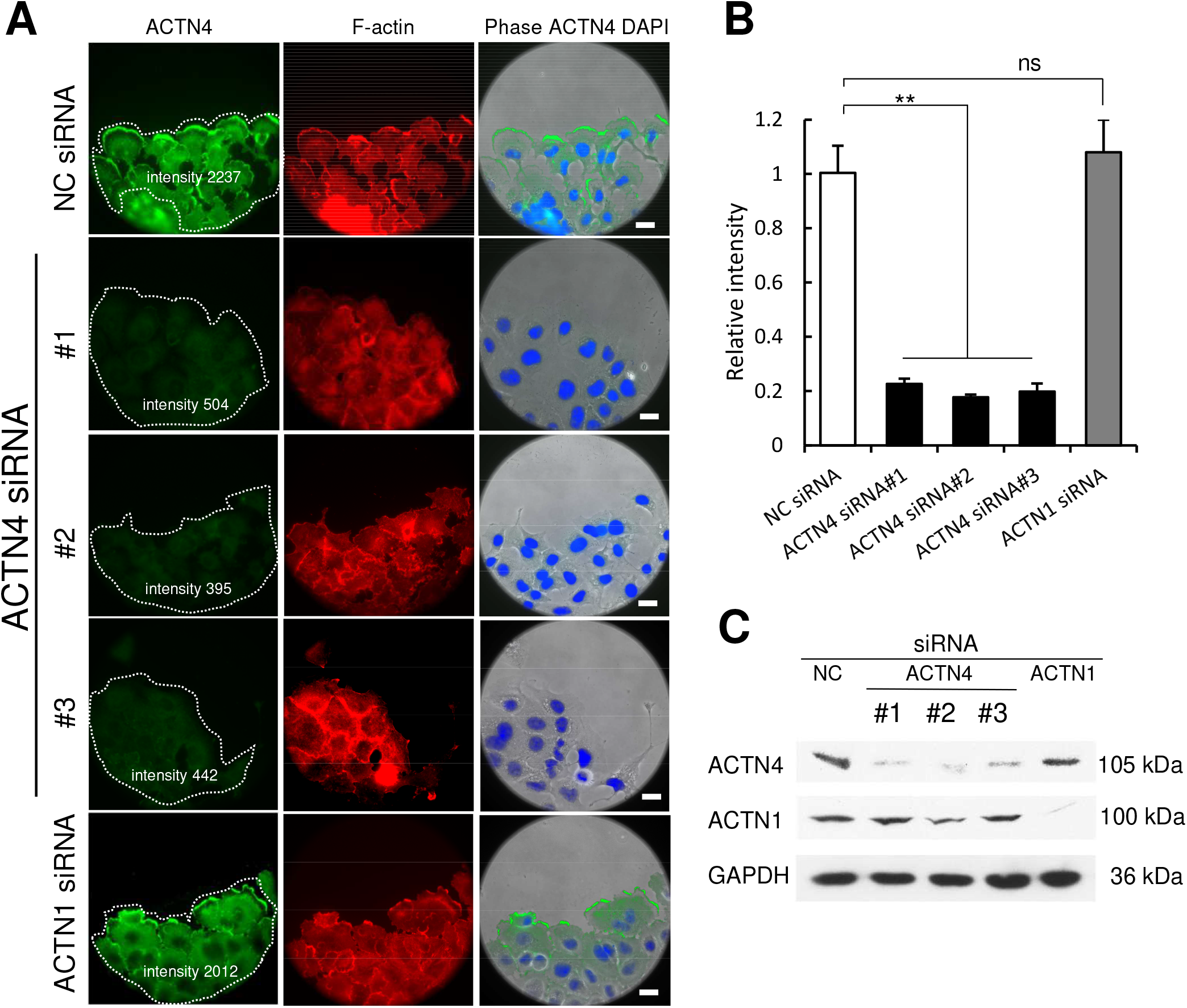
Isoform-specific knockdown of ACTN4 by siRNA. (**A**) A549 cells subconfluently cultured on coverslips were transfected with negative control (NC) siRNA, three ACTN4 siRNAs (#1, #2, #3), or ACTN1 siRNA. Five hours after scratching the monolayer of culture cells, the cells were fixed and subjected to ACTN4 immunocytochemistry and F-actin staining. Nuclei were stained with DAPI. Scale bars: 20 μm. (**B**) Expression levels of ACTN4 were quantified using immunofluorescence intensities measured by the MetaMorph imaging software. The average fluorescence intensity values in >8 cell areas facing the wounded space in each coverslip were calculated. ACTN4 fluorescence intensity was normalized to the intensity of NC siRNA-transfected cells after background correction. ACTN4 siRNA (#1-3), but not ACTN1 siRNA, significantly reduced ACTN4 protein expression (**p<0.01, n=3). Values are expressed as the means ± SEM of triplicate experiments and are representative of two other independent experiments. The fluorescent intensity in representative images was shown in the cell area (dotted line in A). (**C**) Immunoblot analysis with anti-ACTN4 or anti-ACTN1 antibodies demonstrated that ACTN4 siRNAs #1-3 reduce ACTN4 protein levels, but not ACTN1 protein levels. Immunoblotting with an anti-GAPDH antibody verified equal amounts of total protein loading.

